# Fighting *Aspergillus* infection using biocontrol bacteria: A proof-of-concept of environmental interference in a translational setting

**DOI:** 10.1101/2025.03.27.645790

**Authors:** Fabio Palmieri, Aurélien Trompette, Apiha Shanmuganathan, Aureline Bouchard, Julie Pernot, Thomas Junier, Julia M. Kelliher, Patrick S.G. Chain, Jennifer Foster Harris, Armelle Vallat, Marco Pagni, Christophe von Garnier, Janick D. Stucki, Nina Hobi, Angela Koutsokera, Samuel Neuenschwander, Niki D. Ubags, Pilar Junier

## Abstract

*Aspergillus* fungi are opportunistic pathogens that affect millions of people worldwide. Aspergilli produce organic acids to optimize the environmental pH and match the needs of their enzymatic machinery. In this study, we tested the hypothesis that this also occurs during infection. By producing oxalic acid (OA), *Aspergillus* would manipulate pH during lung infection and thus, interfering with this process could control the pathogen. To test this hypothesis, we assessed *in silico* the potential for OA production in a wide range of Aspergilli. A genetic marker for AO production was detected in most of the species including prevalent human pathogens. We tested OA production *in vitro* in four strains of *A. niger* and *A. fumigatus*, but only one of the *A. niger* strains produced OA consistently. For this fungal strain, oxalotrophic bacteria were able to control fungal growth via OA consumption. To translate this observation into a pre-clinical system, increasingly complex experiments were performed. In 3D-cell cultures, *A. niger* also secreted OA and modified pH and free Ca^2+^. Co-inoculation of the oxalotrophic bacterium inhibited the development of the fungus. However, biocontrol could not be replicated in *Galleria mellonella*, which is often used as an infection model. In contrast, the bacterium improved disease score and the absence of oxalate crystals in the lungs in the mouse model. This biocontrol interaction between oxalotrophic bacteria and *A. niger* represents a paradigm shift in the fight against opportunistic fungal pathogens, where the goal is to render the host environment less permissive to pathogen development

**One Sentence Summary:** Demonstration of biocontrol as a therapeutic concept to combat *Aspergillus niger* with oxalotrophic bacteria in an animal infection model

## INTRODUCTION

Recent estimates indicate that 3.8 million people worldwide die due to a fungal infection each year and more than one billion people are estimated to live with a chronic fungal disease (*1*). These numbers are considered conservative, as surveillance and monitoring of fungal burden in human health are limited (*1*). The situation is expected to worsen with the worldwide rise of antifungal resistance in both medical and agricultural settings, and the emergence of novel pathogenic species driven by climate change (*2*). This is notably the case for infections caused by fungi of the *Aspergillus* genus. *Aspergillus* spp. are opportunistic pathogens that affect 14 million people worldwide (*3*). Aspergillosis is manifested in diverse ways ranging from colonization (i.e., presence of fungi without clinical symptoms or radiological or laboratory signs of active disease) to infection (chronic aspergillosis, invasive aspergillosis). These manifestations depend largely on the immune status of the patient (*4–7*). For instance, invasive pulmonary aspergillosis (IPA), is the most severe and lethal form, only occurring in immunosuppressed individuals. The incidence of IPA is approximately 250’000 cases annually and the mortality rate ranges from 30 to 80% (*8*). Given its relevance to public health, *Aspergillus fumigatus,* the most common causative agent of aspergillosis, has been recently classified as one of the four fungal pathogens of critical importance in the first fungal priority pathogen list published by the world health organization to guide research and public actions (*9*).

Aspergilli are widespread in the environment and produce a high quantity of airborne conidia, which can reach densities of up to 10^8^ conidia per m^3^ of air (*6, 10*). In addition to its ubiquity and pathogenic potential, the rising of antifungal resistance in *A. fumigatus* is a challenge in clinical management (*11*). The overuse of antifungal agents, particularly azoles, both in medical and agricultural settings, has led to increasing cases of resistance, considerably limiting treatment options (*11, 12*). Notably, a recent study revealed that many azole-resistant *A. fumigatus* clinical isolates had pre-acquired resistance from the environment, underscoring the impact of environmental exposure to these antifungals (*13*). Moreover, Morogovsky et al. (*14*) demonstrated that azole resistance can also be transmitted between fungal strains via horizontal gene transfer, highlighting the potential for rapid dissemination of resistance among *A. fumigatus* populations. In this context, there is an urgent need for the development of new therapeutic options and innovative solutions to tackle this major threat and to improve patient outcomes (*15*).

Considering that disease manifestations do not result only from the presence of the pathogen or a permissive host, but also from multifactorial environmental conditions that may favor the development of the pathogen, focusing on the environment constitutes a novel treatment approach. In this way, the arena of disease emergence, which includes the host’s environment and its inhabitants (i.e., microbiota), could be the focus of therapeutic interventions (**Fig. 1A**) (*7*). It is now accepted that the microbiome contributes to human health and that changes in the composition or the relative abundance of its constituents can lead to disease (*16*). Accordingly, influencing the microbiome with the intention to maintain or improve health is becoming an alternative treatment approach, and several microbiome-based therapies are currently in development for this purpose (*17*). Although most studies concerning the respiratory tract are descriptive, recent data provide clear evidence of the role of the airway microbiota on respiratory health (*18*). This control can be exerted, for example, through modulation of the host’s immune response or mucus production (*19*).

**Figure 1.**
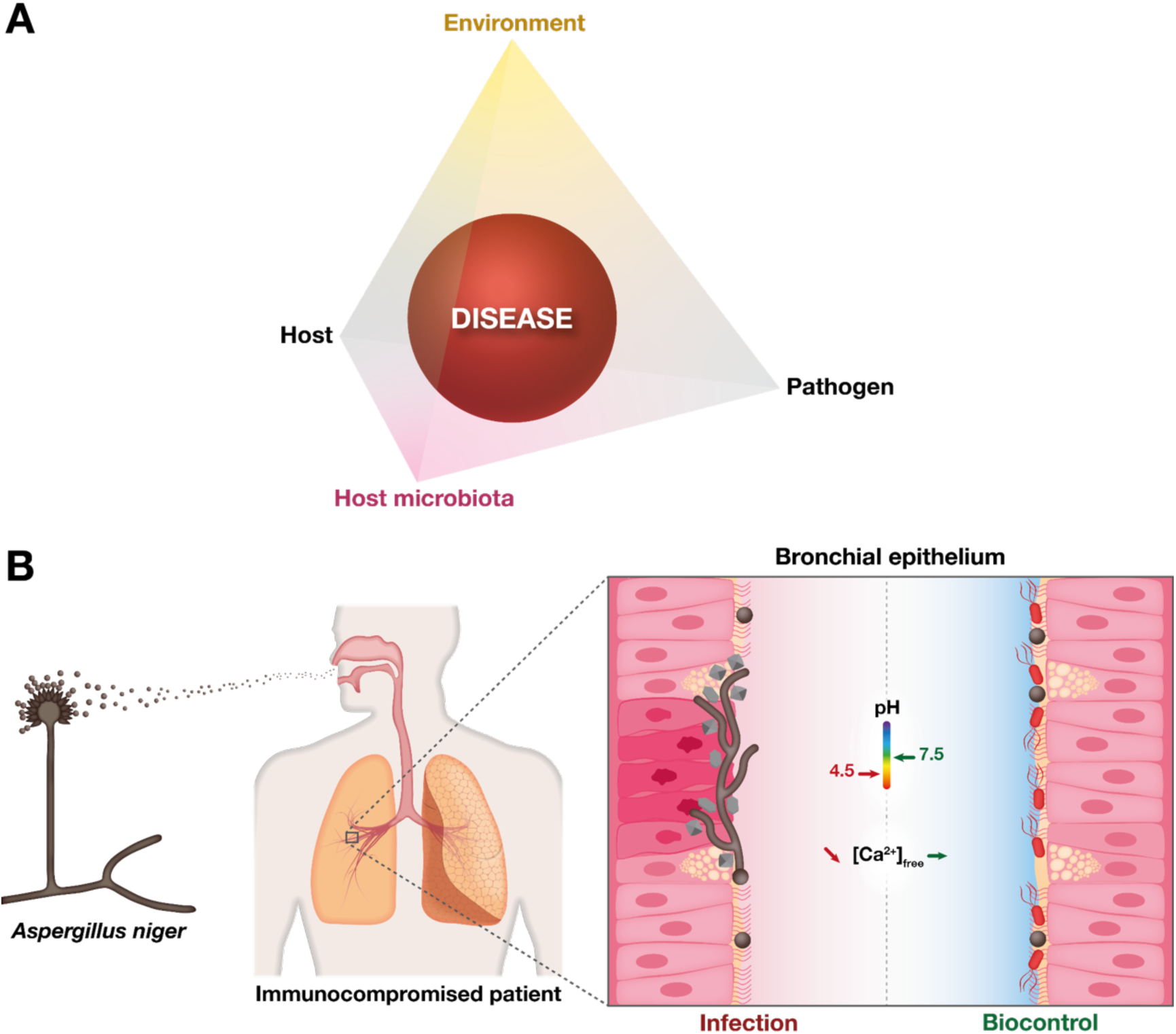
Graphical summary of the environmental interference biological control concept focusing on the consumption of calcium oxalate by oxalotrophic bacteria to inhibit the growth of *Aspergillus niger*. (**A**) A shift in the “host-pathogen” paradigm that considers not only the interaction of the pathogen with the host, but also the role of the resident microbiota and environmental conditions (microbiome) offers a more complex view of disease progression and results in the possibility of focusing on the microbiome to develop novel therapeutic interventions (adapted from (*7*)). (**B**) Example of microbiome manipulation using metabolic processes associated with the production and consumption of oxalic acid (OA). Airborne conidia of *Aspergillus* spp. (depicted in dark brown) arrive in the respiratory tract through breathing. During a normal infection process (left side of the image), *Aspergillus* spp. modifies the environment by secreting OA causing a decrease of pH and chelation of free calcium ions (Ca^2+^) in the form of calcium oxalate (CaOx; depicted in light grey) crystals. This results in the damage of the host’s tissue and inflammation. The microbiome manipulation strategy (right side of the image) takes advantage of the ability of oxalotrophic bacteria (cells depicted in red) to consume CaOx to reestablish physiological pH and free Ca^2+^ concentrations. This results in the biocontrol of the pathogen.

In this study, we aimed to demonstrate the validity of the principle of environmental interference and disease control using the metabolic processes associated with oxalic acid (OA) production and consumption. Several clinical reports have shown the presence of calcium oxalate (CaOx) crystals in pulmonary aspergillosis in animals and humans (*20–25*). Oxalic acid and oxalate crystals are thought to directly cause damage to the host tissues (including pulmonary blood vessels), and to generate free radicals that can harm cells indirectly (*24*). A recent case report of IPA in a 69-year-old man with lymphoma and pneumonia indicated the presence of CaOx crystals around blood vessels and within the blood vessel walls suggesting a potential mechanical role of oxalate crystals in the angioinvasion of *Aspergillus* (*25*). However, a link between OA production and pathogenicity has not yet been demonstrated in fungi from this genus or in any other fungal pathogen in humans. Oxalic acid is a ubiquitous compound in the environment and is thought to have a central role in fungal metabolism (*26*). Its production and consumption by microorganisms have been directly associated with pH regulation in soils (*27*). Thus, the ability of oxalotrophic bacteria to degrade OA produced by *Aspergillus* spp. will result in the control of pH and calcium chelation, which are the two key environmental factors modified by the secretion of OA (**Fig. 1B**). To test this hypothesis, we first assessed *in silico* the potential for OA production in a large diversity of fungal isolates by screening for a gene marker involved in OA production and then validated the production *in vitro* in environmental and clinical strains. The biological control (i.e., biocontrol) exerted by oxalotrophic bacteria was then tested in increasingly complex systems including two complementary 3D-cell cultures systems (submerged cultures, Transwell® inserts and lung-on-a-chip (LoC)), a non-mammalian organismal model to study infection (larvae of *Galleria mellonella*), and finally a preclinical murine infection model. Across these different models, relevant biomarkers (pH and Ca^2+^) were validated. Taken together, these results show a proof-of-concept for environmental interference using biocontrol bacteria that can be potentially used in the fight against respiratory *Aspergillus* infection.

## RESULTS

### Assessment of the potential for oxalic acid production in Aspergilli

*Aspergillus niger* has been extensively studied for its ability to produce OA (*28, 29*), but this trait has been less investigated in other species, including major human pathogens such as *A. fumigatus*. To assess the potential prevalence of OA production among a wide range of Aspergilli, we performed an *in silico* screen based on the presence of the gene encoding the oxaloacetate acetylhydrolase (OAH, EC 3.7.1.1), which catalyzes the conversion of oxaloacetate to OA and carbon dioxide. The primary activity of this enzyme has been associated with OA biogenesis in a wide range of Ascomycota, including plant pathogens (*30*). Moreover, disruption of *oah* resulted in the abolition of OA accumulation in *A. niger* (*31, 32*). The presence of OAH was investigated in 1230 genome assemblies corresponding to over 800 Aspergilli species from diverse origins. OAH was shown to be present across several species and there was no significant difference in prevalence in isolates from specific origins (e.g., comparing clinical, environmental, or veterinary origin; **Fig. 2A**). Nonetheless, the trait is not ubiquitous and in species represented only by clinical isolates, prevalence ranged from 0 to 100%. For instance, in the case of *A. fumigatus*, about 25% of the strains corresponded to strains from a clinical origin. All these strains were predicted to be oxalogenic (**Fig. 2B; Table S1).** These results support the idea of OA as a widespread target of microbiome manipulation by Aspergilli but this requires the validation of OA production before the concept is tested *in vivo*.

**Figure 2.**
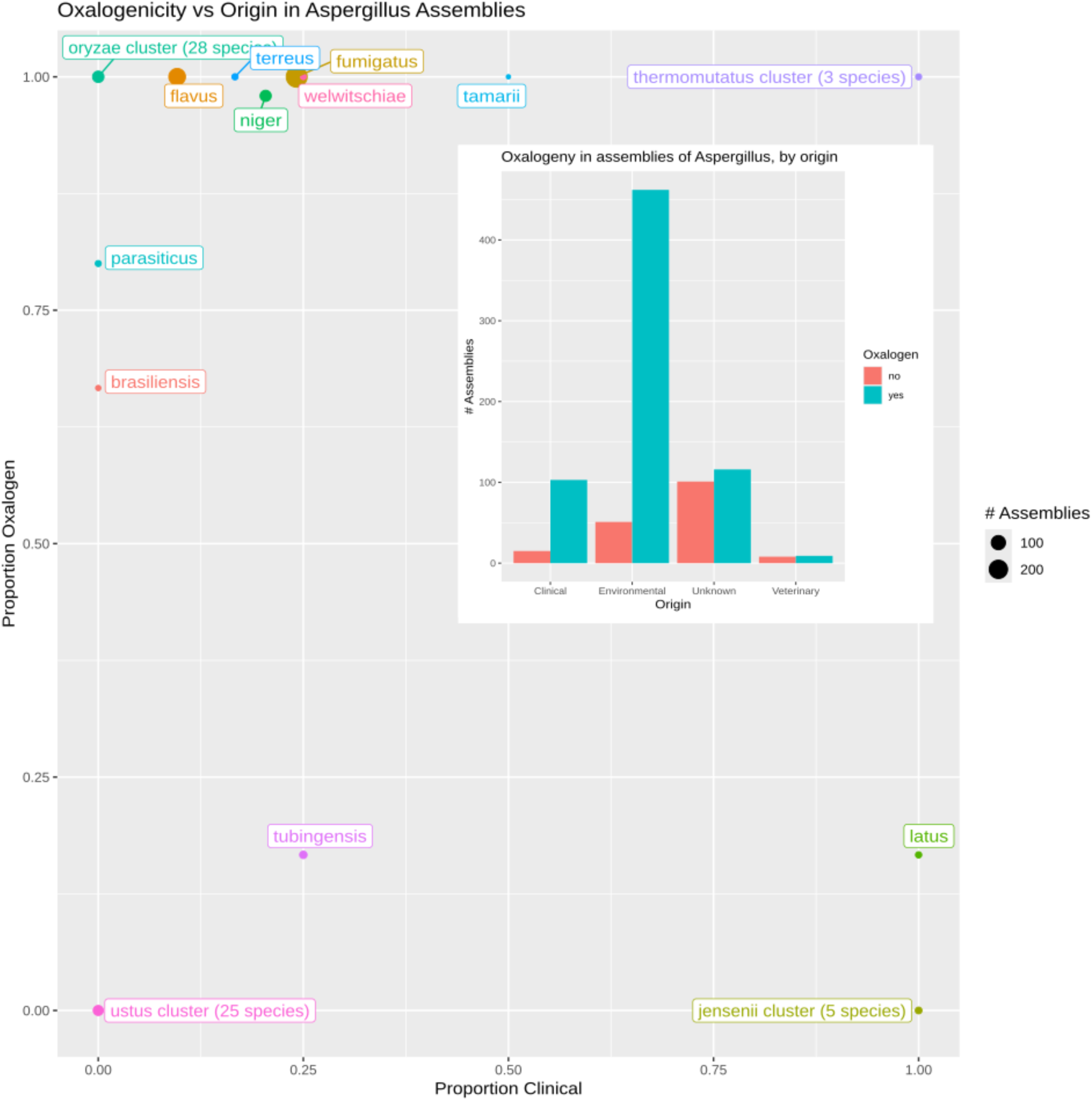
Proportions of oxalogenic *vs.* clinical assemblies of *Aspergillus* within species. Several species are comprised only of oxalogenic assemblies (*e.g. A. fumigatus* (top)), others contain none (*e.g. A. ustus* (bottom)); the same is true regarding clinical origin (*e.g. A. parasiticus* (left) and *A. latus* (right)). As a result, the same position in the scatterplot may be occupied by more than one species. We denote such groups by the term “clusters”. A marker’s size denotes the number of assemblies in the corresponding species; the marker’s color is specific to a position; the label denotes the single species at the given position, or (for clusters) one member of the cluster as well as the number of species within it. Information on the composition of the species clusters is provided in **Table S1**.

### *In vitro* validation of oxalic acid production and biocontrol in environmental and clinical Aspergillus strains

Since the metabolism of fungi can change significantly depending on the nutritional conditions of the growth medium, we evaluated OA production in Aspergilli using different culture media. We tested four fungal strains belonging to *A. niger* and *A. fumigatus*. The two *A. fumigatus* strains (Af293 and CEA10) corresponded to clinical isolates (*33*). The two *A. niger* strains corresponded to a strain used in the industry for citric acid production (A1144) (*34*) and to an environmental strain (M8) from our strain collection. Acidification was first detected in potato dextrose agar (PDA) in the presence of a pH indicator. While the two strains of *A. niger* (M8 and A1144) acidified the medium, only one of the strains of *A. fumigatus* (CEA10) acidified the medium (**Fig. 3A)**. Acidification could be the result of OA production, but also of the production of other organic acids such as citric acid (*28, 35*). Therefore, Ultra-High-Performance Liquid Chromatography (UHPLC) was performed in supernatants from liquid cultures in three different media. This analysis revealed that strain M8 is the only one consistently producing OA regardless of the medium. Two of the three clinical isolates (*A. niger* A1144 and *A. fumigatus* CE10) produced OA in two of the media, while *A. fumigatus* Af293 only produced a marginal amount in R2 medium (**Fig. 3B**). We thus concluded that *A. niger* M8 consistently produced OA and acidified the pH under laboratory conditions and selected this strain for further experiments. Next, we tested the effect of confrontation of the fungus with *Cupriavidus oxalaticus*, an oxalotrophic bacterium (*36, 37*). The confrontation was performed using the Air-Liquid Interface (ALI) medium, which is a medium commonly used for differentiation of lung cells (*38*). *Cupriavidus oxalaticus* was found not only to control growth, but also to stabilize the pH of the culture medium at a neutral pH (**Fig. 3C**). This was consistent with a decrease in OA detected in the medium (**Fig. 3D**). We therefore concluded that oxalotrophic bacteria have a significant inhibitory effect on *A. niger* OA production and growth *in vitro*.

**Figure 3.**
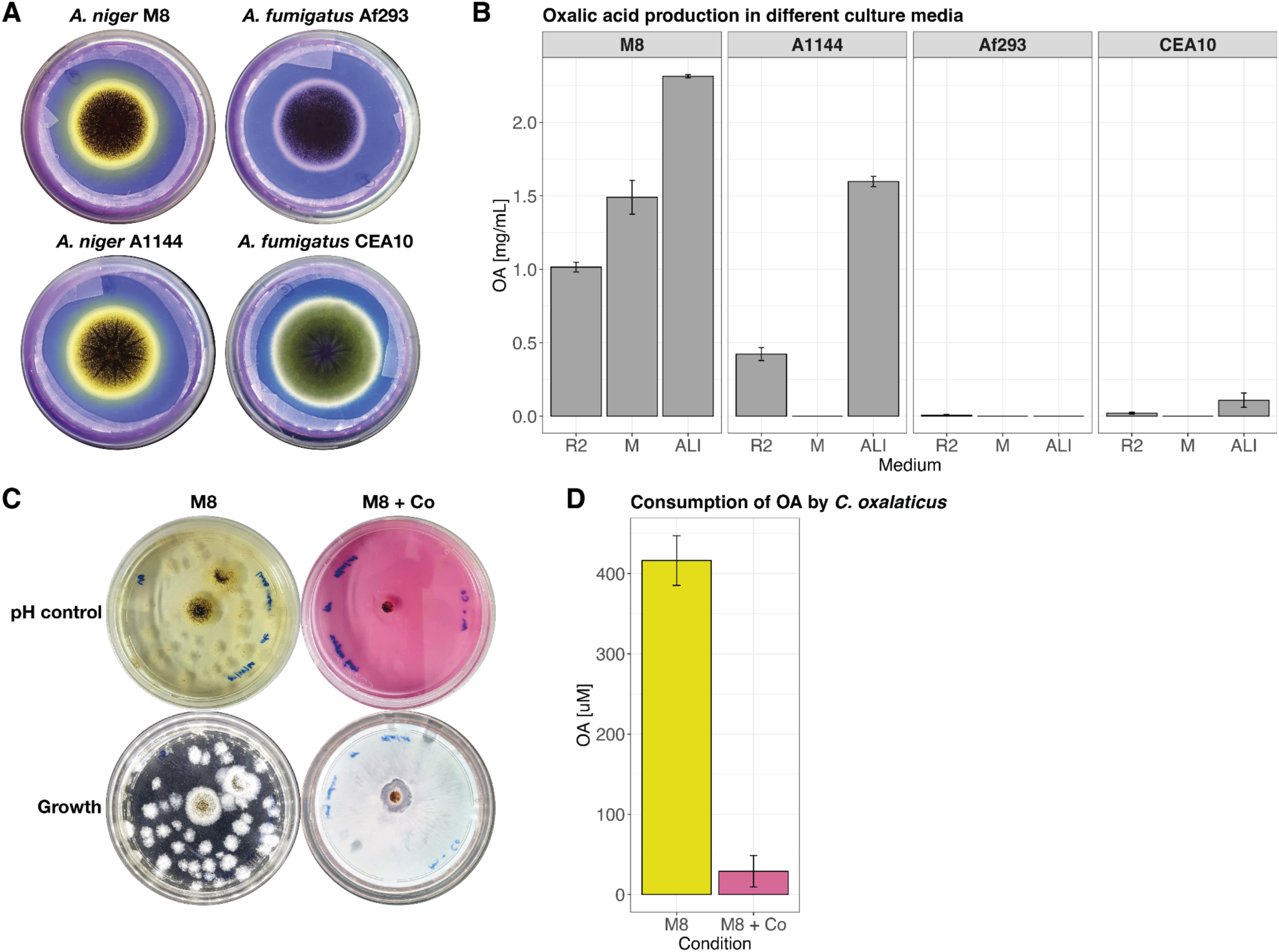
Experimental validation of the production of oxalic acid (OA) in Aspergilli in different culture media and biocontrol through oxalotrophy. (**A**) Acidification test in Potato Dextrose Agar media supplemented with 50 mg/L bromophenol blue (BPB). The tests were performed with two strains from *A. niger* and *A. fumigatus*, respectively. Three of the strains correspond to clinical isolates (A1144, CE10, and Af293). Both *A. niger* (M8 and A1144) acidify the medium, as shown by the change in the color of the agar from purple blue to yellow. In contrast, only *A. fumigatus* CEA10 only slightly acidifies the medium, while no change was observed for *A. fumigatus* Af293. (**B**) Quantification of OA by UHPLC. *A. niger* M8 is the only strain constitutively producing OA in the different liquid media tested. R2 = Reasoner’s 2A medium; M = Malt broth; ALI = Air-Liquid Interface medium. *A. fumigatus* CEA10 produced OA in the ALI medium and marginally in R2 medium. (**C**) Confrontation assay between *A. niger* M8 (M8) and the oxalotrophic bacterium *C. oxalaticus* (Co) in ALI medium. The top row shows the difference in pH (yellow for acidic pH and pink for neutral) for both treatments: fungus alone (M8) and fungus in confrontation with the bacterium (M8 + Co). The bottom row shows images of the same plates but using a light source to better visualize the growth of the fungus in both conditions. (**D**) Oxalic acid concentration decreased by close to 90% in presence of the bacterium (M8 + Co) as compared to the control with the fungus alone (M8).

### Validation of physiological readouts in submerged bronchial cell cultures

An animal-free human bronchial epithelial cell (HBEC) submerged culture lung infection model compatible with the 3R principles of animal experimentation (*39*) was set up to confirm the physiological readouts and a dose-response curve was performed to identify the optimal cell concentration and relative ratios of *A. niger* and *C. oxalaticus* to be used in follow-up experiments. The overall size, shape, and integrity of the lung cells was monitored after 24 h. In the co-cultures with *A. niger* the three parameters changed at an absolute load of 500 conidia and above. The HBECs shrank in size due to actin agglomeration. Moreover, from a conidial load above or equal to 1000 conidia, the fungus also had an adverse effect on tissue integrity, and hyphae were observed in the cultures (**Fig. S1**). The same experiment was performed with *C. oxalaticus*. A morphological change was observed in cell load equal and above 500 bacterial cells. The HBECs co-cultured with *C. oxalaticus* became rounder, and actin agglomeration increased compared with the cells-only control (**Fig. S2**). Finally, we analyzed the effect of co-culturing *A. niger* with the oxalotrophic bacterium on HBECs integrity. The experiments were performed with 10 or 500 conidia and 10 bacterial cells. After 72h, *A. niger* induced morphological changes (size reduction and actin agglomeration), with a stronger effect for 500 conidia, confirming the results obtained at 24 h. With the co-inoculation of as few as 10 *C. oxalaticus* cells, the morphology of the HBECs was like the morphology of HBECs in the bacteria-only control, suggesting the inhibition of fungal development. Moreover, hyphae were not observed in the co-cultures (**Fig. S3**). We concluded that a conidial and bacterial load of 10 conidia/cells was ideal to monitor the interaction of *A. niger* and *C. oxalaticus* in differentiated cell cultures.

### *In vitro* biocontrol in bronchial tissue in Transwell® inserts

After establishing a dose-response curve on HBECs in submerged cultures, the effect of inoculation of 10 *A. niger* conidia alone or in co-culture with 10 *C. oxalaticus* cells was assessed in differentiated bronchial tissue in Transwell® inserts. A strong cytopathic effect of *A. niger* was observed based on the integrity of the monolayer of differentiated bronchial epithelial cells. In the presence of the fungus, the cell monolayer is strongly damaged compared to cells incubated without the fungus (**Fig. 4A)**. Moreover, the presence of the fungus induced changes in pH and Ca^2+^concentration, the two environmental factors predicted to vary upon the production of OA. The pH dropped significantly from 7.5 down to 4.5 (**Fig. 4B**). This was directly correlated with a strong decrease in Ca^2+^ concentrations from 1 mM to around 0.2 mM (**Fig. 4B**). In contrast, pH and Ca^2+^ levels were statistically indistinguishable when oxalotrophic bacteria were co-cultured with the fungus and compared to the controls with bronchial cells alone or with bacteria. In addition, CaOx crystals were observed in the cultures in which the fungus developed, but not when the fungus was co-cultured with oxalotrophic bacteria (**Fig. 4C**). This suggests that the lower levels of Ca^2+^ measured in the treatment with the fungus were likely the result of complexation with OA and corroborates with the decrease in pH. Moreover, the absence of CaOx crystals when the fungus was in co-culture with *C. oxalaticus* agrees with the pH and Ca^2+^ concentration measured.

**Figure 4.**
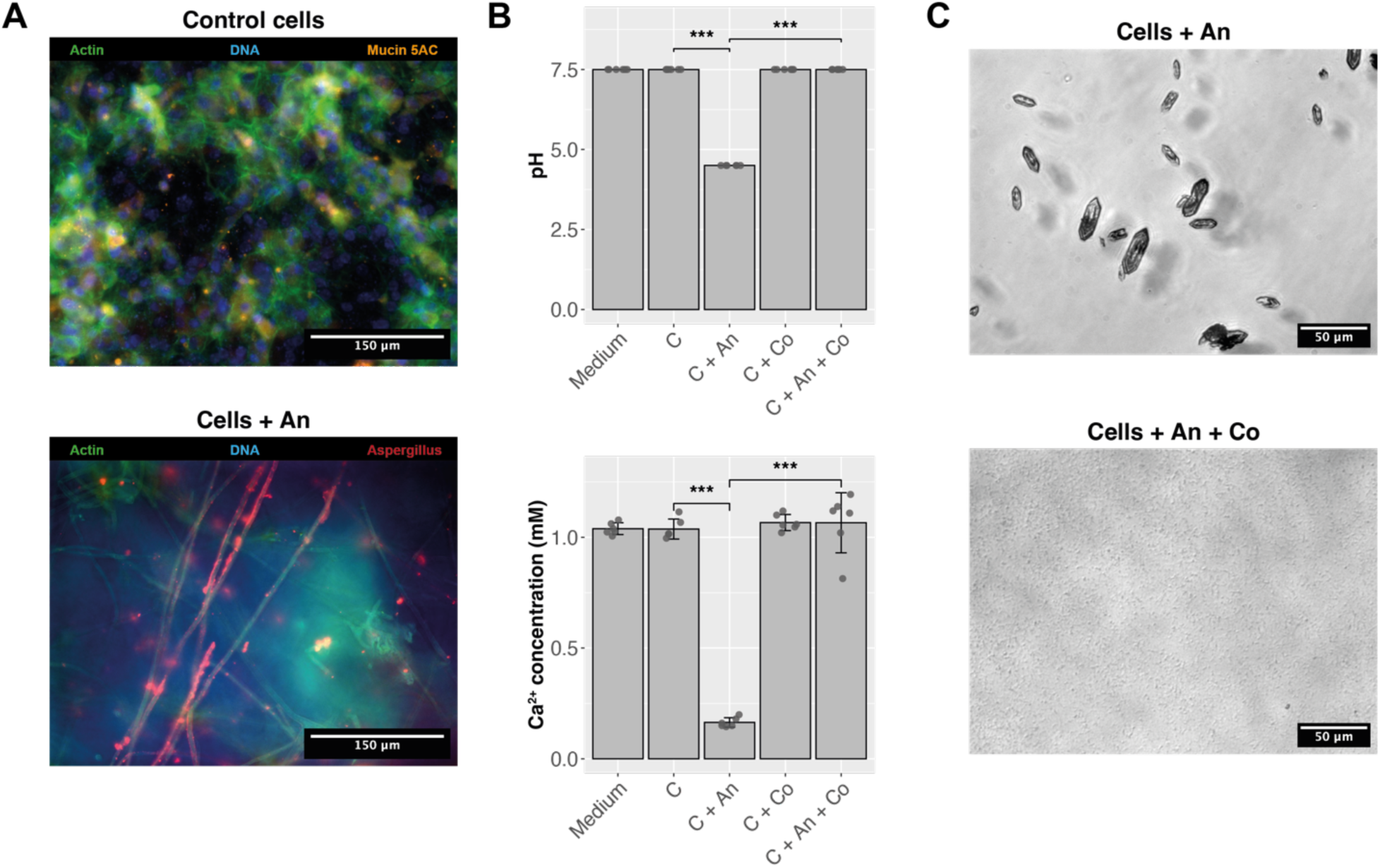
Validation of physiological readouts in differentiated bronchial tissue in Transwell® inserts. **(A)** Immunofluorescence microscopic pictures showing healthy control differentiated bronchial epithelial cells (upper panel, Control cells), and damaged bronchial epithelial cells infected with *A. niger* (lower panel, Cells + An) after 72 h of incubation. **(B)** pH and calcium (Ca^2+^) concentration measurements in the different treatments. In the presence of the fungus (An), the pH decreases, as compared to all other treatments. This pH decrease is correlated with a drastic decrease in the concentration of free Ca^2+^. However, in the co-culture with the oxalotrophic bacterium (Co), pH and free Ca^2+^ concentration return to physiological levels. ***p < 0.001. **(C)** CaOx crystals in the presence of the fungus (Cells + An, upper panel), and absence of CaOx crystals in coculture with oxalotrophic bacteria (Cells + An + Co, lower panel). An = *A. niger*, Co = *C. oxalaticus*.

### Validation of OA production by *A. niger* and biocontrol on chip

To assess whether *A. niger* exposure in an *in vitro* culture would lead to infection and associated OA production in alveoli, we exposed immortalized alveolar epithelial cell line (AXiAECs) (*40*) seeded on an AX12 LoC under static and submerged conditions. Epithelial barrier function was measured repeatedly, and we observed a decrease in TEER starting after 26 h post-exposure (**Fig. 5A**) Interestingly, we observed CaOx crystal formation in the culture media at 49 h post-exposure (**Fig. 5B**), which coincided with a restructuring of the alveolar epithelial cells around the area where the crystals were observed (**Fig. 5C**). *Aspergillus niger* infection led to a decrease in pH of the culture medium (**Fig. 5D**) and we also observed a decrease in free Ca^2+^ (**Fig. 5E**), as expected. Prolonged exposure to *A. niger* led to a complete destruction of the alveolar epithelial cells at 72 h post-exposure (**Fig. 5F**) and microscopic examination of the membranes revealed the formation of CaOx crystals (**Fig. 5G**). Taken together, we established a *A. niger* infection model in alveolar epithelial cells cultured on a LoC, which reflects a reduction in pH and free Ca^2+^ and the formation of CaOx crystals. Next, we assessed whether *C. oxalaticus* could rescue the *A. niger*-induced impairment of the alveolar epithelial barrier in the *in vitro* LoC model. Immortalized primary human alveolar epithelial cells were infected with *A. niger* and exposed to either vehicle control or *C. oxalaticus* after 24 h. Exposure to *A. niger* drastically decreased TEER measurements and thus an enhanced impairment of the alveolar epithelial barrier function (**Fig. 6A**). Interestingly, confrontation exposure with *A. niger* and *C. oxalaticus* rescued the observed phenotype and no reduction in TEER was observed. The vehicle control cells had a slightly enhanced TEER, which can be explained by repetitive TEER measurements (**Fig. 6A**). Immunofluorescent imaging of the membranes indicated a reduction in loss of structure which was observed with *A. niger* infection alone (**Fig. S4)**. However, we did not observe any differences in pH or free Ca^2+^ levels between the *A. niger* and *A. niger* with *C. oxalaticus* exposure groups (**Fig. 6C**). Taken together, we demonstrate the feasibility for biocontrol in *in vitro* culture with alveolar epithelial cells.

**Figure 5.**
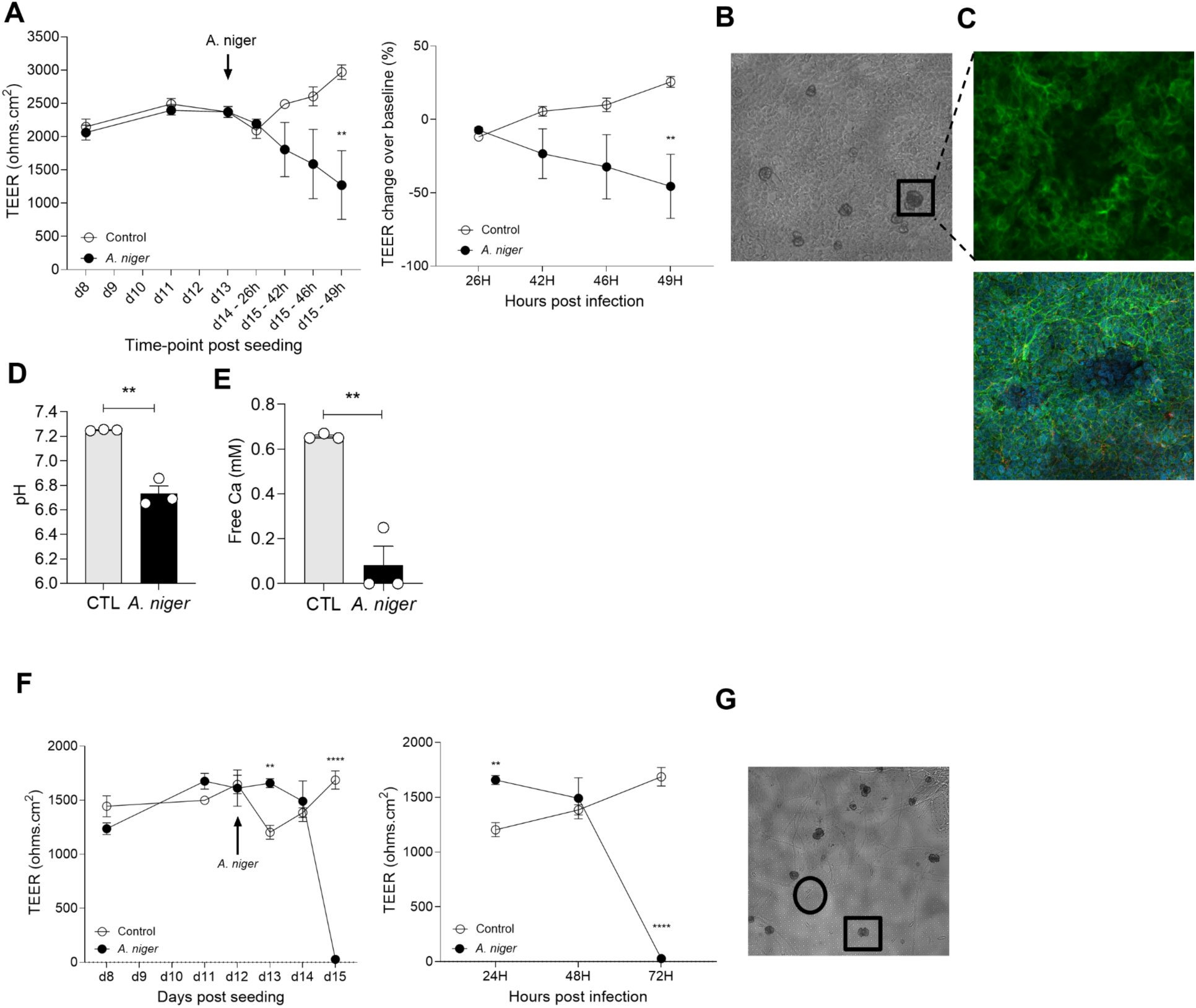
*Aspergillus niger* exposure impairs alveolar epithelial cell barrier. (**A**) TransEpithelial Electrical Resistance (TEER) measurements (indicative of epithelial barrier integrity) in immortalized human primary airway epithelial cells (AECs) following infection with *Aspergillus niger* (10 conidia) up until 49 hours post-infection. (**B**) Presence of calcium oxalate crystals in the culture medium 48 hours post *A. niger* infection. (scale = 20 µm) (**C**) Same area of the inlet in (**B**) showing the degradation of the epithelial monolayer by the crystal agglomerates. (scale = 25 µm and 100 µm) (**D**) Quantification of pH in the culture medium of AECs 48 hours post*-A. niger* infection. (**E**) Quantification of free calcium in the culture medium of AECs 48 hours post*-A. niger* infection. (**F**) TEER measurements in AECs following infection with *Aspergillus niger* (10 conidia) up to 72 hours post-infection showing complete destruction of the epithelial barrier integrity. (**G**) Presence of hyphae and amorphous (as indicated by a square) or fully formed (indicated by a circle) calcium oxalate crystals in the culture medium 72 hours *post-Aspergillus niger* infection.

**Figure 6.**
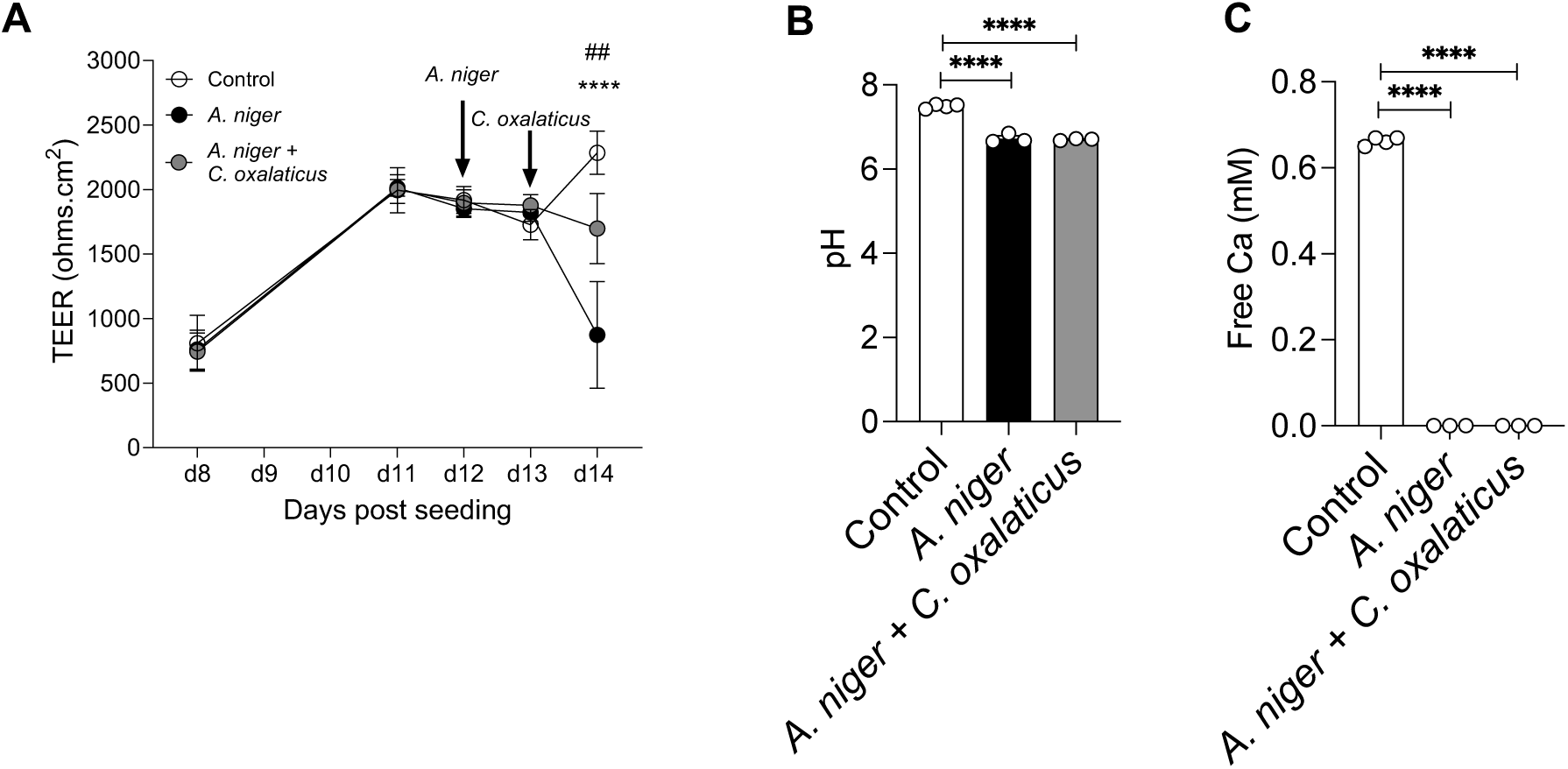
Exposure of alveolar epithelial cells to *C. oxalaticus* rescues *A. niger* induced barrier impairment. (A) *In vitro* model of the “environmental interference concept” confronting the oxalotrophic bacteria *Cupriavidus oxalaticus with Aspergillus niger* and looking at TEER alterations in AECs until 48 hours post-fungal infection. (B-C) Quantification of pH (C) and free calcium (D) in the culture medium of AECs 48 hours *post-Aspergillus niger* infection.

### Tolerance and safety concerns of using a soil bacterium in an animal lung model

*Cupriavidus oxalaticus* is a bacterium that has not been identified in the human or murine lung. Hence, there is a need to determine whether the bacterium itself can be harmful. Preliminary tests with cell cultures showed that *C. oxalaticus* can trigger inflammation (**Fig. S5**) or be cytotoxic (**Fig. S6**). Therefore, we assessed the effect of *C. oxalaticus* exposure on immortalized primary alveolar epithelial cells *in vitro* and in mouse lungs *in vivo*. Immortalized primary alveolar epithelial cells seeded on AX12 LoC in submerged condition were exposed to different doses of *C. oxalaticus* (50, 200, or 800 CFUs). The highest doses had a significant positive effect on TEER measurements at 72 h post-exposure compared to controls (**Fig. 7A**), which is suggestive of enhanced alveolar epithelial barrier function, reflected also by rearrangement of actin (**Fig. 7B**). In addition, immunosuppressed mice were exposed to different doses of *C. oxalaticus* (from 10^2^ to 10^4^ CFUs) via intranasal administration at experimental day 0 **(Fig. 8A**). The different doses of *C. oxalaticus* were well tolerated by the immunosuppressed mice as indicated by a low disease score (**Fig. 8B**). Moreover, no differences were observed in total and differential cell counts in the bronchoalveolar lavage fluid (BALF) of the mice at 24 h post-exposure (**Fig. 8C**), suggesting that *C. oxalaticus* exposure induces limited inflammation in the airways and is thus well tolerated by immunosuppressed mice.

**Figure 7.**
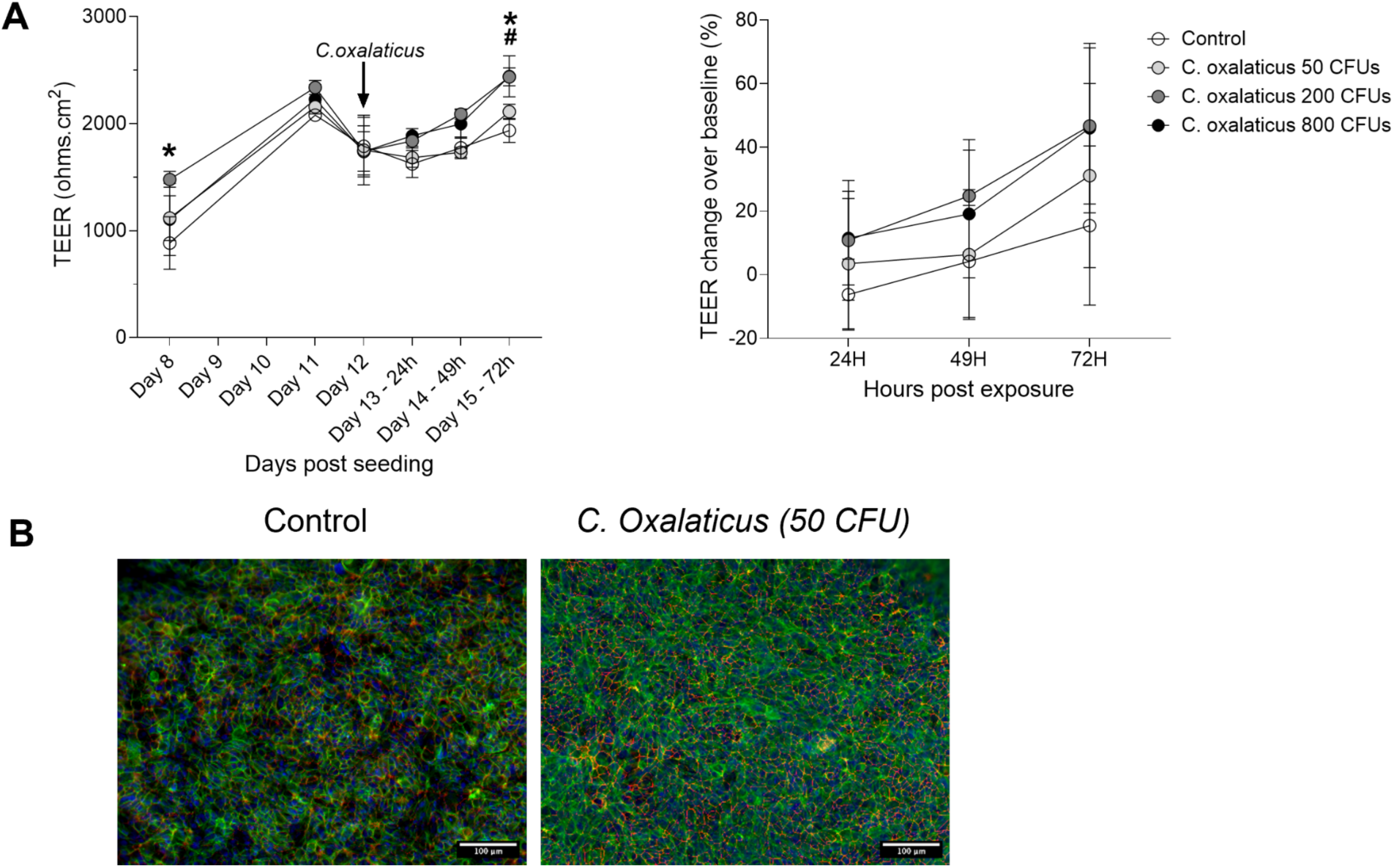
*Cupriavidus oxalaticus* improves alveolar epithelial barrier function. (**A**) TransEpithelial Electrical Resistance (TEER) measurements (indicative of epithelial barrier integrity) in immortalized human primary airway epithelial cells (AECs) following exposure to *C. oxalaticus* (50, 200, or 800 colony forming units (CFUs). (**B**) Representative immunofluorescence microscopy pictures of AECs (magnification 20x, scale = 100 µm).

**Figure 8.**
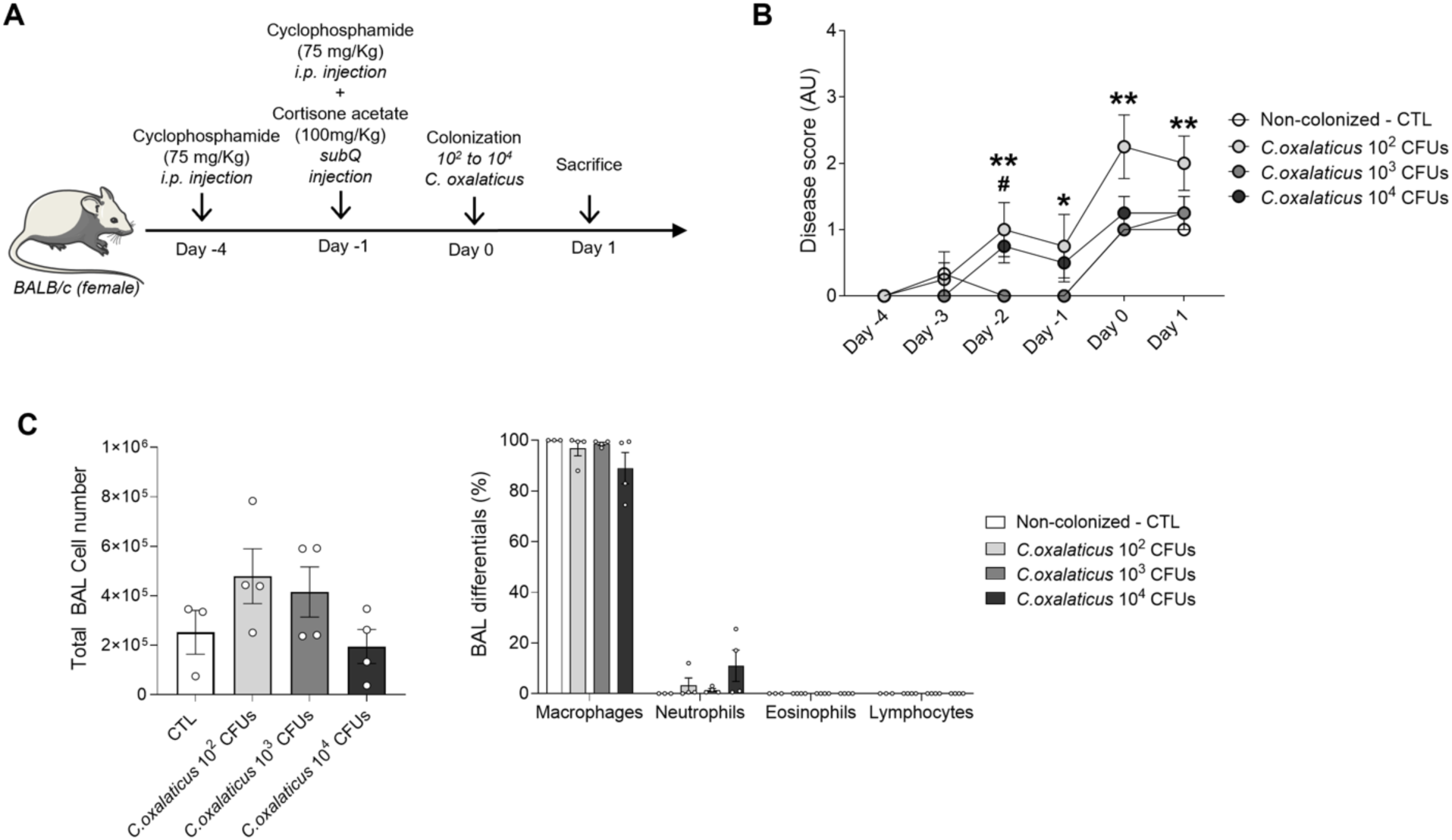
*Cupriavidus oxalaticus* exposure is well tolerated by immunosuppressed mice. **A.** Experimental model of fungal doses titration of *C. oxalaticus* doses (from 10^2^ to 10^4^ CFUs) in immunosuppressed mice with two injection of 75 mg/Kg cyclophosphamide (Day -4 and Day -1) and one injection of 100 mg/Kg cortisone acetate (CA) (Day-1). **B.** Severity assessment after bacterial colonization. Statistics significance were indicated with the following symbols when comparing the control group (Non-colonized CTL) with: (*) *C.oxalaticus* 10^2^ CFUs and (#) *C.oxalaticus* 10^4^ CFUs. C. Cell count and differentials of immune cells (percentage) in the broncho-alveolar lavage fluid (BAL). All data are expressed as mean ± SEM (n = 3 CTL and n = 4 per group). *P < 0.05, **P ≤ 0.01, ***P ≤ 0.001, ****P ≤ 0.0001.

### In vivo biocontrol in Galleria mellonella

To test the biocontrol potential of *C. oxalaticus* against *A. niger in vivo*, we first used *G. mellonella* larvae as a non-mammalian infection model widely used for fungal pathogens (*41, 42*). We started by validating the use of *G. mellonella* larvae for fungal-bacterial co-infection models by repeating a previously published study using *Candida albicans*-*Staphylococcus aureus* co-infections (*43*). We injected 10^5^ *C. albicans* and 2 x 10^4^ *S. aureus*, both alone or as a co-injection, and monitored survival of *Galleria* larvae for 72 h. Survival analysis revealed that both *C. albicans* mono-injections and *C. albicans*-*S. aureus* co-injections were lethal before 24 hours post-infection (hpi, **Fig. S7**), while mono-injections of *S. aureus* were well tolerated by the larvae, with 75% survival after 72 hpi (**Fig. S7**). This is in contrast with the study from Sheehan et al. (*43*), which reported that mono-injections of both *C. albicans* and *S. aureus* were well tolerated by *Galleria* larvae. We next performed a survival assay by injecting 10^7^ *A. niger* conidia and 10^6^ *C. oxalaticus* cells, both alone and in co-injections. We also used heat-killed (HK) *A. niger* conidia and *C. oxalaticus* cells as controls. The injection of 10^7^ *A. niger* conidia led to 100% death of the larvae before 48 hpi. However, injection of 10^7^ HK *A. niger* conidia led to 75% survival after 72 hpi. Injection of 10^6^ *C. oxalaticus* cells, both alive and HK, was well tolerated by the larvae, with 100% and approximately 90% survival after 72 hpi for live and HK *C. oxalaticus,* respectively. In contrast, co-injections of *A. niger* with *C. oxalaticus* (live and HK) were lethal with 100% death before 36 hpi (**Fig. 9A**). Interestingly, the presence of melanized nodules, which is a hallmark of innate immune response in this model (*44, 45*), were noticeable in *A. niger*-infected larvae. To check for hyphal growth, the larvae were dissected and melanized nodules were resected from the haemocoel of larvae. Hyphae were stained using calcofluor white (**Fig. 9B**). A high density of fungal hyphae was visible in larvae injected with *A. niger* conidia. However, in the heat-killed *A. niger* conidia control, no hyphae were present. When *C. oxalaticus* was co-injected with *A. niger*, hyphae were still visible. Finally, we also checked for hyphal and bacterial growth (**Fig. 9C**). We could confirm hyphal growth in the larvae injected with *A. niger* conidia. Moreover, no growth was observed in the case of heat-killed *A. niger* conidia. For the larvae co-injected with *A. niger* and *C. oxalaticus*, both hyphal and bacterial growth were observed.

**Figure 9.**
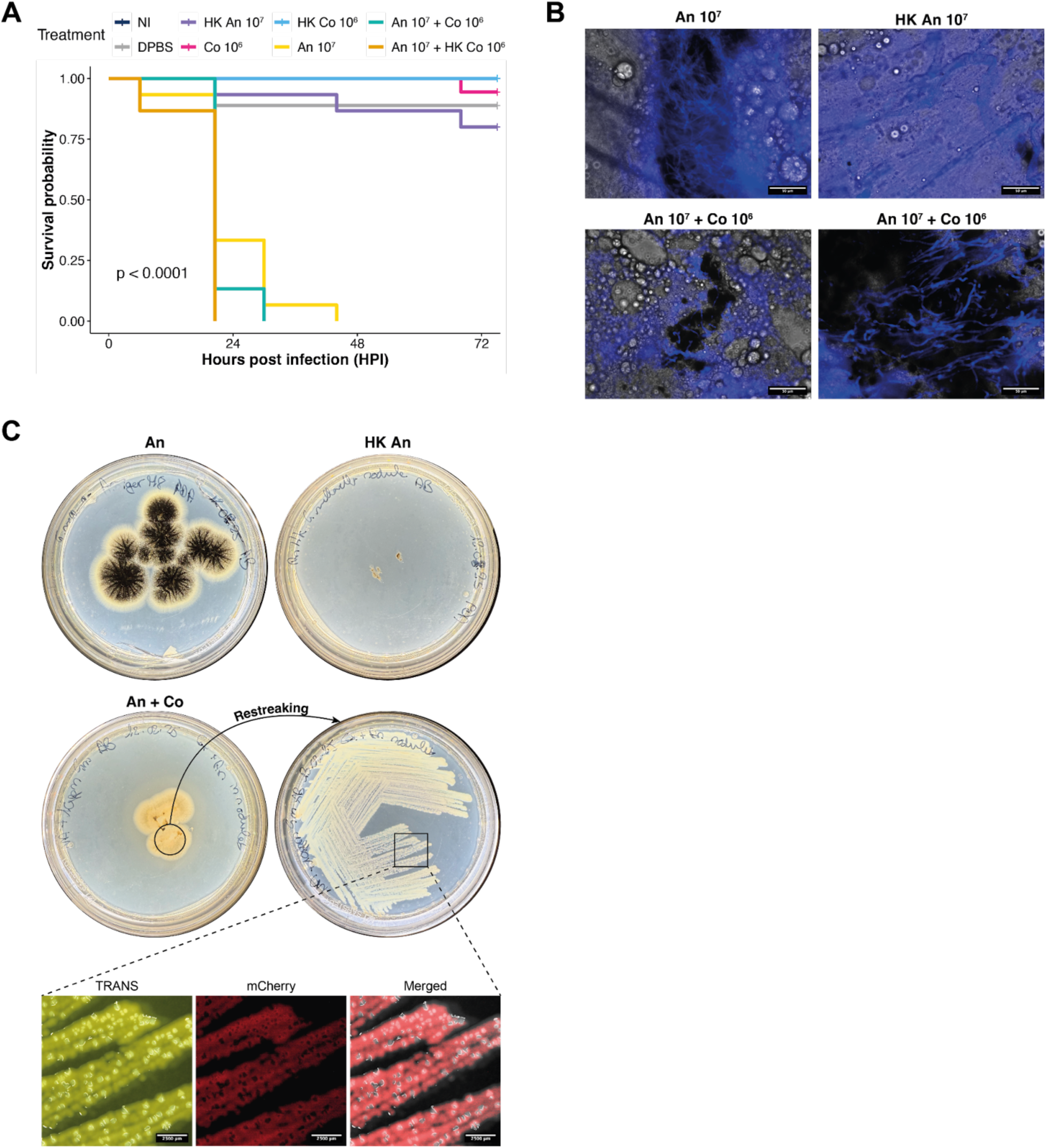
Non-mammalian *in-vivo* biocontrol model in *Galleria mellonella*. **(A)** Kaplan-Meier survival plot of *G. mellonella* larvae injected with *A. niger* (An) and *C. oxalaticus* (Co), alone or in combination. **(B)** Fluorescence imaging of *G. mellonella* nodules stained for fungal hyphae with calcofluor white. **(C)** Images of culture plates to check for fungal growth (PDA), and bacterial growth (NA supplemented with 10 ppm gentamicin) from dissected nodules. Close-up images were taken using an epifluorescence stereomicroscope. NI = no injection, DPBS = vehicle control, HK = heat-killed.

### *In vivo* biocontrol in mice

Until now, no experimental animal models of respiratory *A. niger* infection exist. We developed a murine model of *A. niger* infection in immunosuppressed animals using pre-germinated *A. niger* conidia/hyphae (**Fig. 10A**). Mice infected using different doses of *A. niger* (from 5x10^5^ to 2x10^6^ pre-germinated conidia), had a high increase in disease score at 24h post-infection (**Fig. 10B**). In line with this, we observed hyphae formation in lung tissue sections of these mice (**Fig. 10C**), where the number of hyphae increased with an increasing dose of *A. niger*. Interestingly, we observed the formation of calcium oxalate crystals in areas of the lung with hyphae formation (**Fig. 10D**), which confirms that the experimental murine *A. niger* infection model exhibits specific characteristics required for our biocontrol model.

**Figure 10.**
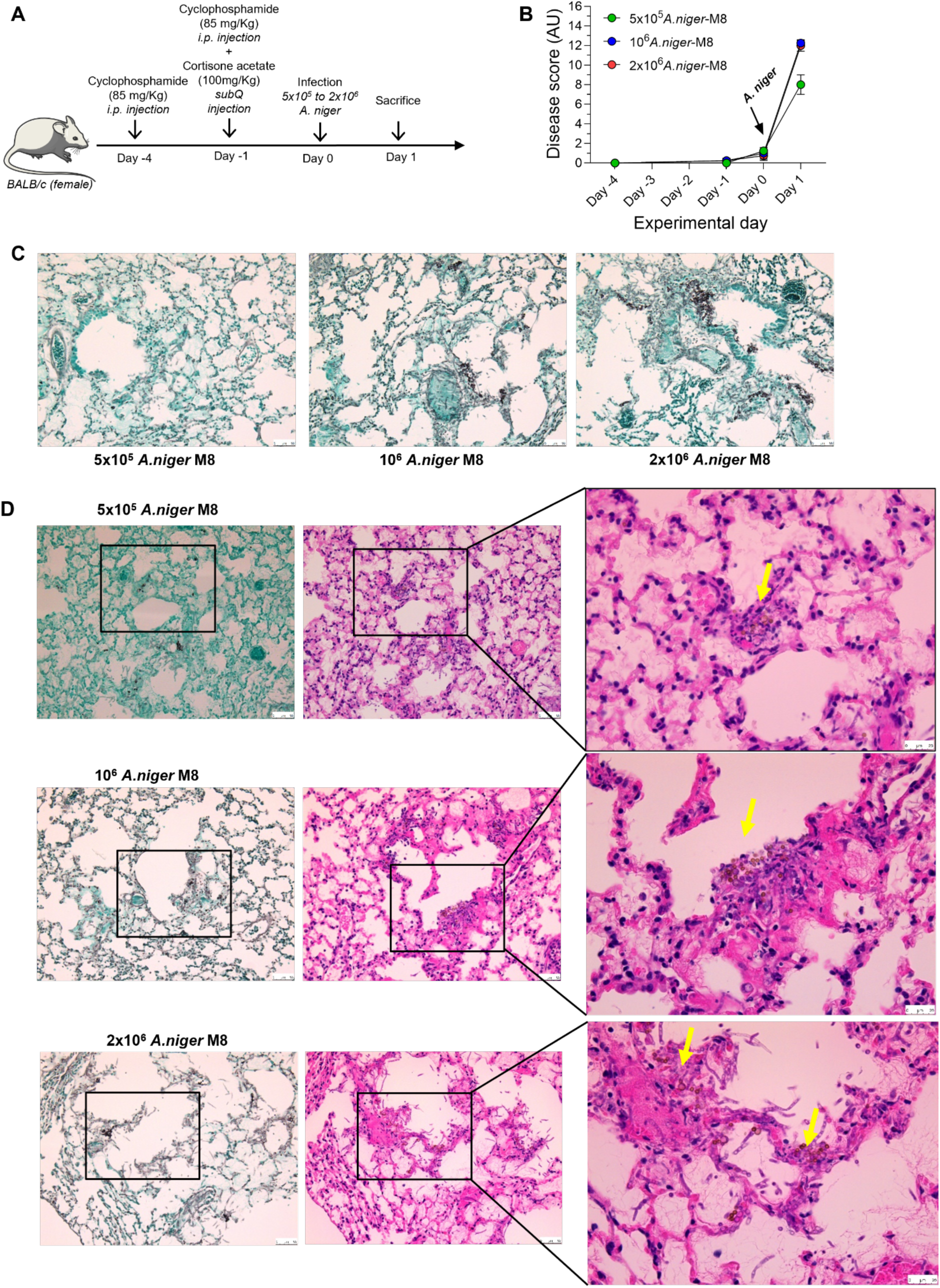
Murine model of respiratory *A. niger* infection. **A.** Experimental model of *A. niger* M8 infection (from 5x10^5^ to 2x10^6^ pre-germinated conidia) in immunosuppressed mice with two injection of 85mg/Kg cyclophosphamide (Day -4 and Day -1) and one injection of 100mg/Kg cortisone acetate (CA) (Day-1). **B.** Disease score assessment after fungal infection. **C.** Representative Grocott’s methenamine silver stain (GMS)-stained of lung tissue from infected mice with 5x10^5^, 10^6^ and 2x10^6^ *A.niger* M8 conidiae (magnification 20x, scale = 50 µm) **D.** Representative GMS and H&E-stained lung tissue sections from infected mice with 5x10^5^, 10^6^ and 2x10^6^ *A.niger* M8 conidiae. Inserts represent areas with crystal formation as indicated by yellow arrows (magnification 20x, scale = 50 µm). All data are expressed as mean ± SEM (n = 4 CTL and n = 4 per group).

To assess whether *C. oxalaticus* exposure could rescue the severity of the *A. niger* infection in our murine infection model, we performed a confrontation experiment. Immunosuppressed mice were exposed intranasally to *A. niger* with or without *C. oxalaticus* (**Fig. 11A**). Mice exposed to both *A. niger* and *C. oxalaticus* had a decrease in disease score compared to mice who were exposed to *A. niger* alone (**Fig. 11B**), suggesting that *C. oxalaticus* can control the severity of *A. niger* infection in an experimental animal model. Although we did not see any differences in total and differential cell counts in the BALF (**Fig. 11C**), we observed a decrease in hyphae formation in the lungs of mice who received both *A. niger* and *C. oxalaticus.* Taken together, these results show a proof-of-concept experimental animal model for biocontrol bacteria in the fight against respiratory *Aspergillus* infection.

**Figure 11.**
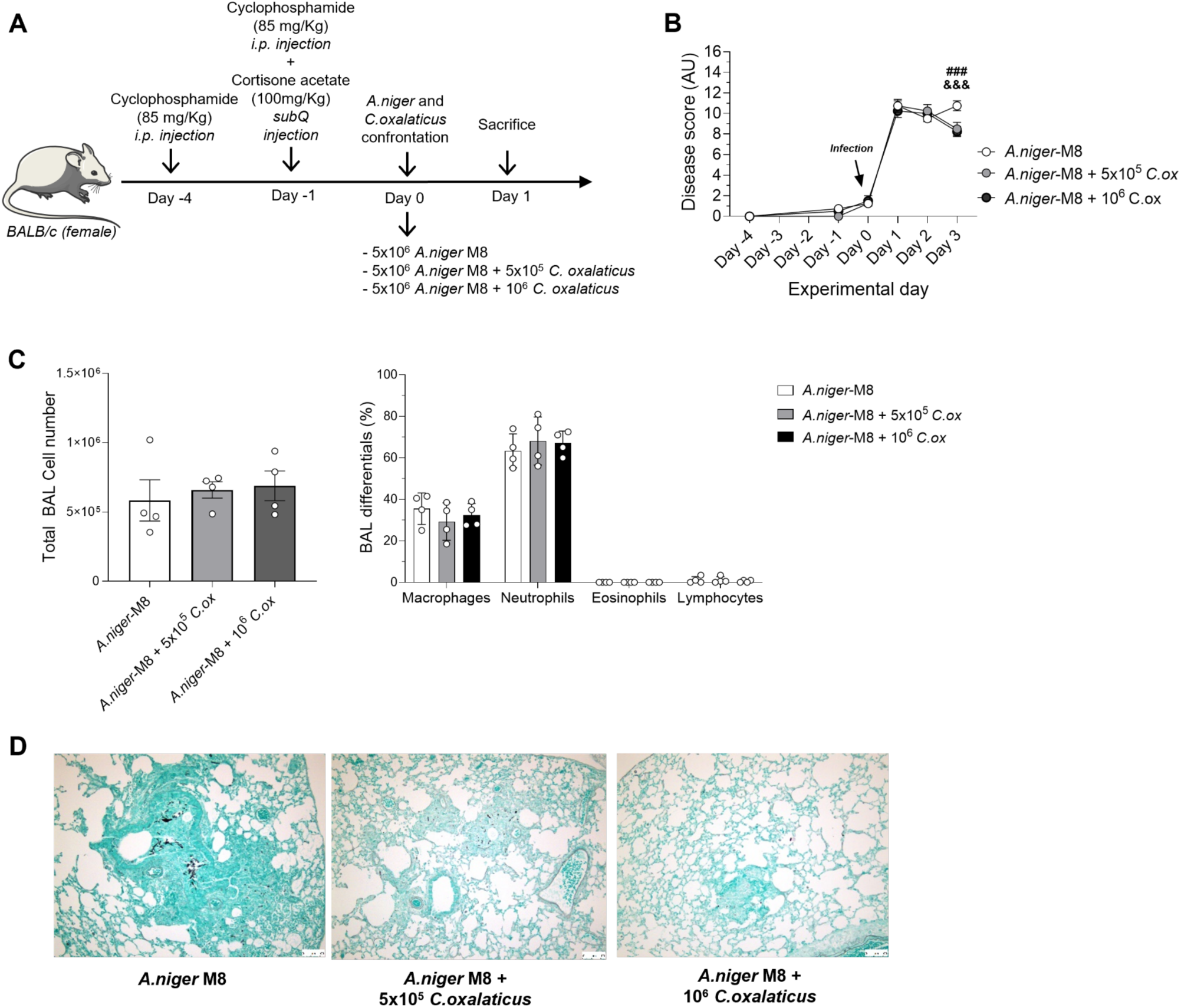
*C. oxalaticus* exposure reduces disease score following *A. niger* infection in mice. **A.** Experimental model of *A.niger M8* (5x10^5^ conidia) and C.oxalaticus (5x10^5^ and 10^6^ CFUs) confrontation in immunosuppressed mice with two injection of 85mg/Kg cyclophosphamide (Day -4 and Day -1) and one injection of 100mg/Kg cortisone acetate (CA) (Day-1). **B.** Disease score assessment after fungal and bacterial confrontation. Statistics significance were indicated with the following symbols when comparing group only infected with with 5x10^5^ *A.niger* M8 conidia: (#) *A.niger* + *C.oxalaticus 5x*10^5^ CFUs and (&) *A.niger* + *C.oxalaticus* 10^6^ CFUs. **C.** Cell count and differentials of immune cells (percentage) in the broncho-alveolar lavage fluid (BAL). **D.** Representative Grocott’s methenamine silver stain (GMS)-stained lung tissue from infected mice with 5x10^5^, 10^6^ and 2x10^6^ *A.niger* M8 (magnification 20x, scale = 50 µm). All data are expressed as mean ± SEM (n = 4 CTL and n = 4 per group). All data are expressed as mean ± SEM (n = 4 CTL and n = 4 per group). Statistics were determined with Two-way ANOVA test (no corrections for multiple comparisons) for **B**, Student’s t-test (unpaired, two-tailed) in C (for BAL cell counts), or multiple t-test (no corrections for multiple comparisons) in C (for BAL immune cells differentials). *P < 0.05, **P ≤ 0.01, ***P ≤ 0.001, ****P ≤ 0.0001.

## DISCUSSION

The growing understanding of the role of the microbiome in human health provides a powerful incentive to consider the microbiome as part of an integrative approach in disease management (*16*). The best examples in which the usefulness of microbiome-based treatments has been demonstrated correspond to diseases affecting the gut, where fecal microbiota transplantation has emerged as a valid and successful alternative to control the expansion of multidrug-resistant organisms or other persistent pathogens such as *Clostridioides difficile* (*46, 47*). Similarly, the concept of nasal microbiota transplantation has been recently proposed as a therapeutic approach to treat chronic rhinosinusitis (*48*). This shows a clear potential for extending the therapeutic use of the microbiota to the respiratory tract. Respiratory tract infections are ubiquitous in society and place a high burden on healthcare systems (*49*). Moreover, multiple studies support the communication between microbiota of the respiratory tract and gastrointestinal tract through the circulatory and immune system (*50*). However, both systems are not only anatomically distinct, but also differ in the abundance and composition of their microbiota (*51*). Therefore, the impact of microbiota modification for therapeutic purposes in lungs need specific considerations and translational efforts. Nonetheless, the lung microbiota can be considered as an unexploited and understudied therapeutic target that may be more prone to modification than other sources of patient heterogeneity, such as host genomes or comorbidities (*52*).

In this study, we validated a novel concept for microbiome manipulation based on the interference with environmental conditions that are permissive for the development of the pathogen. We targeted consumption of OA, a metabolite required for fungal pathogenesis. We demonstrated that the secretion of OA by *A. niger* results in the modification of key environmental parameters during infection in increasingly complex models for aspergillosis. Oxalic acid production has been widely acknowledged as a pathogenicity factor in fungal phytopathogens, such as *Sclerotinia sclerotiorum* and *Botrytis cinerea* (*53, 54*) where pH manipulation and Ca^2+^ chelation plays a direct role in pathogenesis (*26, 55*). Both local acidification and cation complexation is exploited by these plant pathogens to weaken the cell wall structure, facilitate infection, inhibit plant defenses, and induce programmed cell death (*56*). In Transwell® and a LoC model, we have shown that OA secretion resulted in CaOx crystal production, acidification and chelation of free Ca^2+^. The detection of CaOx crystal coincided with remodeling around alveolar epithelial cells, supporting the hypothesis of host tissue damage (*24*). Also, by developing a new murine model of aspergillosis for *A. niger*, we were able to show that CaOx crystal formation is also a biomarker for infection, as suspected by the reported observation of crystals in cases of pulmonary aspergillosis in animals and humans (*20–25*). Overall, our results across multiple models show for the first time that fungal production of OA is relevant in the context of pathogenesis in an animal system.

We showed that oxalotrophic bacteria can be used for the biocontrol of the pathogen, both *in vitro* and *in vivo*. Oxalotrophic bacteria are known inhabitants of the human gut, where they perform the key function of degrading dietary oxalate (*57*). These group of bacteria have also been used as probiotics for the treatment of hyperoxaluria (high oxalate in urine) and the management of kidney stones (*57, 58*). However, they have not yet been proposed for the control of infectious diseases. While oxalotrophic bacteria are well characterized in the gut, this is not the case of the lung. Therefore, we used a soil bacterium for the validation of the biocontrol strategy evaluated here. There are three elements in which any new therapy would be judged during regulatory assessment: quality, safety, and efficacy. Responding adequately to these is the challenge faced by many microbiome-based therapies (*16*). In our case, the use of a soil oxalotrophic bacterium could represent a safety issue, given the immunosuppressed context of the host in which the pathogen develops. Nevertheless, we have shown that the bacterium improves alveolar epithelial barrier function and is well tolerated by immunosuppressed mice, despite the potential pro-inflammatory or cytotoxic effects detected in cell cultures. The high tolerance could be the indirect result of the immunosuppressing treatment, which likely reduced the inflammatory response in the mouse. An altered immune response to parasites has been reported in immunosuppressed mice (*59*), and therefore, additional tolerability assays should be performed with immunocompetent mouse. Nevertheless our results show the value of exploiting a combination of 2D and 3D cell cultures and LoC systems during the validation phase, to assess key pitfalls in the safety assessment during the validation process. A similar approach could be considered when evaluating novel strains to assess the potential impact of bacteria on the integrity of the host epithelial barrier. This is a stage required for considering the broad reprogramming of respiratory mucosal homeostasis, which further involves controlled immune monitoring.

Despite the positive results obtained across most of the models investigated here, the results obtained with *G. mellonella* show some of the limitations of translation. Compared to expensive and difficult to manage vertebrate models, larvae of *G. mellonella* have emerged as a faster, cheaper, and higher throughput model to study fungal infection (*44*). However, one of the areas that has been much less explored is the effect of co-infections. Although we were able to replicate results from a previous co-infection study (*43*) and that we confirmed pathogenesis of *A. niger* and tolerance of *C. oxalaticus* individually, in the co-infection the bacterium did not have a biocontrol effect, contrary to the results obtained in cell cultures, LoC, or the mouse model. This shows a significant limitation of using a non-mammalian model for the fast screening of candidate biocontrol bacteria.

In conclusion, in this study we have demonstrated biocontrol across multiple systems and the translation of the biocontrol concept from a purely *in silico* prediction, to an *in vivo* animal model. In particular, the results obtained with the animal model suggest the therapeutic potential of oxalotrophic bacteria for the control of aspergillosis. This opens the possibility of attempting to harness the lung microbiota for therapeutic purposes. The composition of the lung microbiome is known to be highly dynamic and be modified under the influence of different kinds of diseases. Accordingly, monitoring the presence and changes in the composition of the pulmonary microbiome is a new direction for studying pathogenesis and disease progression (*60, 61*). Oxalate-degrading capabilities have been previously reported in strains of the genera *Lactobacillus* (*62*), *Streptococcus* (*63*), *Prevotella* (*64, 65*), and *Veillonella* (*65*), all of which are reported as components of the lung microbiota. In the future, human-derived oxalotrophic bacteria could be tested to validate a preventive or therapeutic effect over fungal infection.

## MATERIALS AND METHODS

### Bioinformatic analysis

A list of all available *Aspergillus* assemblies was downloaded as ‘ncbi_dataset.tsv‘ from NCBI’s Genome Portal (https://www.ncbi.nlm.nih.gov/datasets/genome/?taxon=5052). Genomic sequences for all assemblies listed in that file were downloaded using the datasets’ tool (*66*) (total number: 1230). When an assembly came in both RefSeq and GenBank versions, only the RefSeq version was kept. All assemblies were subjected to gene prediction using Augustus (*67*). The protein products of the predicted genes were scanned for OAH with the ICL_PEPM (AC: cd00377) model from the Conserved Domains Database (*68*) using HMMer [https://hmmer.org]. This yielded 11019 hits. The ICL_PEPM family contains more members than OAH, however, so most HMM hits were not OAHs. True OAHs were identified using phylogenetic analysis, as follows. Hits of the HMM search were clustered using CD-HIT (*69*), yielding 480 clusters. The 480 cluster heads, as well as 45 reference *Aspergillus* OAHs downloaded from UniProtKB (*70*) and ten outgroup sequences (see below) were aligned with MAFFT (*71*), then trimmed with trimAl (*72*) with a gap threshold of 0.8. The outgroup sequences were chosen at random from CDD’s KPHMT-like family (cd06557), which is closely related to (but distinct from) ICL_PEPM. Sequences shorter than 90% of the full width of the alignment were then discarded. A phylogeny was then built from the resulting alignment using IQ-TREE 2 (*73*). The phylogeny contained a clade containing exclusively reference OAHs from UniProtKB or cluster heads. The cluster heads belonging to that clade were therefore considered probable OAHs. CD-HIT cluster members corresponding to cluster heads thus identified as probable OAHs were then extracted (using the Newick Utilities (*74*)), for a total of 1006 probable OAHs. The species to which each assembly belongs was extracted from ‘ncbi_dataset.tsv‘, and each species were mapped to its origin using the ‘isolation_sourcè field returned by ‘datasets summary genome …‘ for each accession in ‘ncbi_datasets.tsv‘. These were manually categorized into clinical, veterinary, environmental, or unknown. Join operations resulted in a 885-line TSV file linking assembly AC, organism name, origin, and oxalogenicity.

### Bacterial and Fungal cultures

All bacterial and fungal strains originated from the culture collection of the Laboratory of Microbiology of the University of Neuchâtel (**Table 1**). *Cupriavidus oxalaticus* Ox1 was tagged in-house using insertion with a MiniTn7 system (*37*). **Table 2** summarizes all the media used. Bacterial strains were routinely cultured on NA medium. *Aspergillus niger* was routinely cultured on MA medium. PDA was used for *A. niger* conidia production.

### LMWOA detection by colorimetric pH indicator-based Petri dish assay and UHPLC

For the culture-based assay, Potato Dextrose Agar (PDA) supplemented with 50 mg/L bromophenol blue (BPB) was used as a pH indicator-containing medium. Culture plates were imaged after two weeks of incubation at room temperature (RT). For the UHPLC analysis, 500 µl of 30 mM H2SO4 were added to 1 mL of a two-week liquid culture in malt 1/10, Reasoner’s 2, and ALI liquid media in triplicate, to obtain 20 mM H2SO4 final concentration to obtain a low pH for the extraction of LMWOAs and to dissolve any precipitated crystals. The samples were incubated at 60°C for two hours to dissolve precipitated metal oxalate crystals, and then centrifuged at 3000 g for 10 min. All the samples were filtered at 0.22 µm (13mm syringe filters, PTFE, hydrophilic) and 200 µl were added into HPLC vials with 250 µl conical inserts. UHPLC (Ultimate 3000 RS-Dionex, Thermo Fisher Scientific, USA) was coupled with a DAD detector set at 210 ± 2 nm. A 5 µL of sample was injected onto a PrevailTM organic acid column (5 µm particle size, 150 x 4.6 mm, Grace Davison Discovery Sciences, USA) with the temperature kept at 40°C. The mobile phase consisted of 50 mM phosphate buffer adjusted to pH 2.5 with phosphoric acid with a flow rate of 1 mL/min. Pure oxalic acid (Merck, Germany) was identified by the retention time and was quantified by an external standard curve, linear regression from five calibration points (0.2 to 5 mg/mL).

### Primary normal human bronchial epithelial cell culture

Primary normal human bronchial epithelial cells (Lifeline Cell Technology, USA) were expanded in a T-75 cell culture flask with vent cap (Corning, USA) in BronchiaLife™ B/T complete medium (Lifeline Cell Technology, USA) supplemented with 0.5% Phenol Red solution (Sigma-Aldrich, USA, 15 mg/L final concentration) to 70-80% confluence in a humidified incubator at 37°C with 5% CO_2_. Culture medium was changed every other day. Cells were used until passage 2 for all experiments. Cells were harvested by trypsinization with 0.05% Trypsin / 0.02% EDTA (Lifeline Cell Technology, USA), followed by the addition of Trypsin Neutralizing buffer (Lifeline Cell Technology, USA), and counted using a hemocytometer after centrifugation at 100xg for 5 min and resuspension of the cell pellet in BronchiaLife™ medium.

Cells were seeded at a density of 3x10^4^ cells/well in 200 μl BronchiaLife™ medium for submerged undifferentiated tissue culture in 96-well plates, and 5x10^4^ cells and 8.6x10^4^ cells in 200 μL BronchiaLife™ medium for air-lifted differentiated tissue culture in the apical side of Transwell® inserts in 24-well plates (Corning, USA) and BoC devices, respectively. Transwell® inserts and BoC were first coated with collagen (30 μg/mL) prior seeding of the cells in order to allow proper cell attachment onto the porous membrane, as described in Arefin et al. (55). 600 μL of BronchiaLife™ medium was added to the basolateral side of Transwell® inserts in the 24-well plates and 3 mL in the basolateral side of the BoC device. 96-well plates, Transwell® inserts, and BoC devices were placed in a humidified incubator at 37°C with 5% CO_2_ for 2-3 days until confluence and formation of a monolayer of bronchial cells. For differentiated bronchial cell tissues (Transwells® and BoCs), cells were shifted to air-liquid interface by removing carefully the BronchiaLife™ medium from the apical side and replacing it by Air-Liquid Interface (ALI) Epithelial Differentiation Medium (Lifeline Cell Technology, USA) supplemented with 0.5% Phenol Red solution (Sigma-Aldrich, USA, 15 mg/L final concentration). The same was done for the medium on the basolateral side. Finally, the medium on the apical side was removed and the inserts and devices were placed in a humidified incubator at 37°C with 5% CO_2_ for 21 days. Medium was changed every other day as described previously. The cultures were observed daily using an EVOS™ XL Core bright field inverted microscope (Thermo Fisher Scientific, USA).

### Determination of conidial and bacterial cell load and confrontation assay on submerged undifferentiated bronchial epithelial cell cultures

To determine the optimal conidial and bacterial load to be used for confrontation on bronchial tissue cultures, increasing conidial and bacterial loads were tested to assess their effect on the morphology of bronchial epithelial cells in submerged cultures. *Aspergillus niger* was cultured on PDA for 7 days at 37°C to produce conidia. Conidia were then harvested as already described. Finally, conidia were resuspended in 2 mL DPBS and quantified with an Improved Neubauer counting chamber. Bacteria were cultured in BronchiaLife™ medium at 37°C overnight and quantified with an Improved Neubauer counting chamber. A stock suspension of *A. niger* conidia was made at 10^6^ conidia/mL that was diluted further to obtain suspensions at 5x10^5^ to 5×10^3^ and 10^3^ conidia/mL. The same was done for *C. oxalaticus* from a stock suspension at 10^5^ bacterial cells/mL diluted until 10^3^ bacterial cells/mL. 10 μL of each suspension was added to submerged undifferentiated bronchial tissue in a 96-well plate to have 10^4^ to 10 conidia/well (200 μL) for *A. niger,* and 10^3^ to 10 bacterial cells/well (200 μL) for *C. oxalaticus*. The plates were placed in a humidified incubator at 37°C with 5% CO_2_ for 24h. For the confrontations assay, cells were infected with 10 or 500 *A. niger* conidia and were put in confrontation with 10 bacterial cells. The plate was placed in a humidified incubator at 37°C with 5% CO_2_ for 72h. After incubation, cells were fixed, stained (actin and nucleus), and imaged as described below (Immunofluorescence staining).

### Immunofluorescence staining

Undifferentiated and differentiated bronchial tissues (Transwell® and BoCs) were fixed with 100 μL 4% paraformaldehyde in DPBS for 15 min at RT. Cells were then rinsed 3 times with 200 μL DPBS, with 2 min waiting time between each rinse. Cells were permeabilized with 100 μL 0.5% Triton X-100 in DPBS for 15 min at RT and rinsed 3 times with 200 μL DPBS, with 2 min waiting time between each rinse. After that, cells were blocked with 100 μL 3% BSA in DPBS for 1h at RT. Anti-Mucin 5AC mouse monoclonal antibody (Abcam, USA, Cat.# ab218466) was prepared in DPBS (1/100). The actin stain (ActinGreen™ 488 ReadyProbes™ Reagent) and the nuclei counterstain (NucBlue™ Live ReadyProbes™ Reagent) were added to the same buffer (2 drops/mL and 1 drop/mL, respectively). Anti-Aspergillus rabbit polyclonal antibody (Abcam, Cat.# ab20419) was also added (1/200) in the staining buffer for the conditions where A. niger conidia were inoculated. Fixed cells were incubated with 100 μL buffer containing the stains and Anti-Mucin 5AC and Anti-Aspergillus antibodies overnight at 4°C. The next day, fixed cells were washed 3 times with DPBS and secondary antibodies were applied. Goat anti-Mouse IgG antibody (1/250) conjugated with Alexa Fluor 546 (Thermo Fisher Scientific, USA, Cat.# A-11003,) directed against Anti-Mucin 5AC antibody and Goat anti-Rabbit IgG antibody (1/500) conjugated with Alexa Fluor 594 (Thermo Fisher Scientific, USA, Cat.# A-11012) directed against Anti-*Aspergillus* antibody were prepared in DPBS. Fixed cells were incubated with 100 μL buffer containing the secondary antibodies overnight at 4°C. The following day, fixed cells were once again washed 3 times with DPBS and the membranes from the inserts were carefully cut out with a sharp knife. The membranes were mounted on a glass slide using Fluoromount-G™ Mounting Medium (Thermo Fisher Scientific, USA) and imaged with a Zeiss Axio Observer Z1 fluorescence inverted microscope (Carl Zeiss AG, Germany).

### pH measurements and quantification of calcium, oxalic acid and lactate dehydrogenase (LDH)

pH measurements of the culture medium after 72h incubation were done directly after taking the samples out of the incubator using pH-indicator strips (Merck, Germany) for the Transwell® inserts samples, and CG8+ i-STAT cartridges for the BoC devices, as the pH change of the culture medium indicator was less visible. Free calcium (Calcium Colorimetric Assay, Sigma-Aldrich, Germany), free oxalic acid (Oxalic Acid Colorimetric Assay Kit), and LDH (ScienCell Research Laboratories, USA) were quantified in the culture medium using colorimetric assay kits following the manufacturer instructions.

### Lung-on-Chip

#### ^AX^Lung-on-chip system

The AXLung-on-chip sytem (AlveoliX, Bern, Switzerland) is composed of Lung-on-chip (AX12). The Lung-on-chip can be connected via a platform AXDock to one electro-pneumatic device, AXExchanger. The plates were provided by AlveoliX (Switzerland) and were seeded with immortalized alveolar epithelial cells (AXiAECs) derived from primary AECs from human lungs (*40*). Media exchange on the AX12 have been performed according to manufacturer’s instructions (AlveoliX, Bern, Switzerland). A volume of 200μL antibiotics mix with streptomycin and penicillin diluted in Alveolar epithelial barrier medium was added in each well (ThermoFisher Scientific, Switzerland) (AlveoliX, Switzerland). Then, the medium with antibiotics was removed via a medium exchange at Day 11. At Day 12 post seeding, different doses of *A. niger* and CFU of *C. oxalaticus* bacteria were prepared in 70μL of media to be put in AX12 wells alone or in confrontation fungi-bacteria.

#### TEER measurements

Transepithelial barrier electrical resistance (TEER) measures were taken at specific time-point using Millicell-ERS2 Volt-Ohm Meter on each well. TEER (W) values were subtracted by the background measurement and multiplied by the area surface of the well (0.071 cm2) to have a final value as W*cm2.

#### Cell fixation and staining

Cells were fixed by a first step of permeabilization with 0.1% Triton, followed by a blocking step with 2% BSA. Then, cells compartments were stained with NucBlue to label the DNA in the nucleus, ActinGreenTM 488 ReadyProbesTM reagent to bind to F-actin and Alexa594-conjugated anti-ZO-1 targeting Zonula-occludens-1 (Zo-1) a tight-junction protein. Immunofluorescence images were obtained on Zeiss Upright Axiovison fluorescence microscope. They were then processed on ImageJ 1.53t software. The Brightness and contrast were adjusted in addition to the color balance.

### Galleria mellonella experimental model

*A. G. mellonella* larvae were reared following a previous study (*75*) from larvae obtained from Royal Pêche (La Chaux-de-Fond, Switzerland). First to sixth instar larvae were housed at 30°C in the dark in plastic boxes equipped with metal mesh lid (NHBS, BugDorm BD5003) filled to the middle heights of the box with their standard diet (100 g wheat flour, 100 g corn flour, 100 g wheat germ, 100 g powdered milk, 75 mL honey, 75 mL glycerin). Humidity was maintained by sprinkling water over the food every 2 days.

After pupation, pupae were placed in individual plastic boxes partially filled with wood shavings, reaching up to the midpoint of each box, and kept at room temperature witha normal day-night cycle.

Once emerged, adults were transferred to plastic containers equipped with a nylon mesh lid (NHBS, BugDorm BD5002). The box was equipped with a parchment paper sheet folded in an accordion shape and held together at the two ends by an aluminum foil. Adults underwent the same treatment as pupae, namely day-night cycle at room temperature. A sex ratio 1:1 for adults was not necessary considering a male can fertilize multiple females. Males and females were differentiated by visual inspection (*75*). We ensured the presence of no more than 10 adults per box. After 7 to 10 days, adults were removed from the plastic boxes and, if not already dead, they were transferred to small insect containers and placed at -20°C. The eggs were retrieved by either placing the paper fold in a bugdorm box with nylon mesh (NHBS, BugDorm BD5002) containing their standard diet or by gently scrubbing the dry egg lining. The rearing boxes were placed at 30°C in the dark and sprayed every day to prevent desiccation. After 10-14 days, the first larvae emerged from the eggs. Temperature at the bottom of the box was monitored every day by touch to assess the overcrowding of the box and, if applicable, the larvae were separated into 2 boxes with fresh diet provided. Approximately 30-40 days after egg-laying, the first fifth instar larvae could be observed. Larvae considered fit for injection (cream-colored with no prior sign of melanization, active (movement without stimulation) and weighing at least 0.3 g) were transferred to a small plastic insect container filled with wood shavings and kept at 10°C for a maximum duration of 3 weeks. A portion of the larvae was kept inside their original rearing box to allow them to finish their life cycle and obtain a new generation.

#### Larvae maintenance and injection

Larvae were kept in the dark at 37°C, 24 h prior to injection. Larvae were starved at least 24h before injection. 10-15 healthy individuals per treatment (no signs of melanization) were selected. Prior to injection, the distal part of the abdomen was disinfected with a cotton swab soaked in 70% ethanol. The larvae were then gently held still and subsequently injected on the last right proleg with either DPBS for control or 10 μL microbial suspension using a Hamilton® 25 µL syringe and a 30Gx Terumo® AGANI needle (AN*3013R1). For coinfection, larvae were injected 1 time on each side, with 10μL of each microbial suspension or 10μL of DPBS (PanBiotech, P04-36500) for the control on each side of the last prolegs. If the assays involved both mono- and co-infection, the mono-infected larvae were injected with DPBS on one side and microbial suspension on the other side. Once the injection received, the larvae were placed in sterile 6-well plates (TC-Platte 6 Well, Suspension F, Starstedt, 83.3920500) and then incubated at 37°C for 72h.

#### Survival analysis

Mortality reports took place 2 times at 9am and 5pm each day during the 72h of the duration of the experiment. Individuals were considered dead if no reaction at all ensued a few touches on different parts of their body. At the end of the experiment, the larvae were sacrificed by being put at -20°C.

*Nodule plating and nodule preparation for microscopy with and without Calcofluor White staining* Larvae from the treatments *A.niger* 10^7^, *A.niger* HK 10^7^, *A.niger* 10^7^ + *C.oxalaticus* 10^6^ and *A.niger* 10^7^ + HK *C.oxalaticus* 10^6^ were chosen at random 24 HPI. The larvae were sacrificed by cutting of the head. The larvae were then opened longitudinally (sagittal plane) with scissors and spread on a petri dish with tweezers. Nodules, recognizable as black spots on the body of the larva, were taken gently with tweezers and put on a microscopic slide. A drop of Calcofluor White was put on the nodule to stain for fungal hyphae and then covered with a cover slip. The slide was left for 1min in the dark before removing the excess liquid with a tissue and subsequent visualization. Additionally, nodules were plated on PDA and NA + 10 ppm Gentamycin after having been gently crushed. They were subsequently left for 48h at 37°C (5% CO2, 100% humidity).

#### Statistical analysis and figures

All statistical analysis were run using R and R studio (version 4.3.2). Data was compiled with Excel (version 2412). Survival analysis to generate Kaplan-Meier plots were made with the packages survival, survminer and ggplot2 from survival tables.

### Experimental animals

Seven-week-old female BALB/c mice were purchased from Charles River Laboratories (L’Arbresle, France) and were housed in a Biosafety level-2 (P2) environment. Animal experiments were done in conformity with institutional guidelines, Swiss federal and cantonal laws on animal protection (Cantonal authorization: VD3668 – VD3668x1).

### Experimental animal models

#### Immunosuppression

Immunosuppression in mice was induced using intraperitoneal (i.p.) administration of cyclophosphamide (75 and 85mg/Kg in sterile Natrium Chloratum 0.9%) (Baxter Healthcare Corporation, Thetford, England) at 4 days and 1 day before infection or colonization combined with subcutaneous (s.c.) administration of cortisone 21-acetate (CA) (100mg/Kg in Dulbbecco’s-PBS) (Sigma-Aldrich, Buchs, Switzerland) 1 day prior to infection or colonization.

#### C. oxalaticus exposure

Immunosuppressed mice were colonized with 10^2^ to 10^4^ CFU of *Cupriavidus oxalaticus* (C.ox.) in 50μL volume via intranasal (i.n.) administration following anesthesia with isoflurane.

#### Aspergillus niger *infection*

Immunosuppressed mice were infected with 10^4^ to 2x10^6^ *A. nige*r M8 conidia (A.n-M8) in 80 μL final volume via intranasal (i.n.) administration following Ketamine (75.75 mg/Kg)/ xylazine (12.08 mg/Kg) anesthesia. *A. niger-M8* conidia were pre-germinated prior to administration. Pre-germination of conidia was performed in a liquid culture medium (Potato Dextrose Broth (PDD)). In brief, *A.niger* M8 conidia suspension (200uL) were cultured in T75 flasks with 50mL of Potato Dextrose Broth (PDD)) and incubated at room temperature for 28 hours. Subsequently, the fungal cultures from the flasks were collected and washed with PBS. The fungal pellet was resuspended in PBS and the number of conidia per mL was counted using a Neubauer chamber. The percentage of sprouting conidia was approximately 15%. Serial dilutions were performed based on the total count of conidia per mL to achive the different fungal doses. For the confrontation experiment, *A. niger* M8 and *C.oxalaticus* were prepared as explained above. The suspension of fungi and bacteria at the indicated concentrations were prepared in 40 μL volume and combined to reach a total volume of 80 μL.

#### Animal wellbeing

The general well-being and behavior of the experimental mice was monitored daily throughout the experiments and based on the Clinical Scoring (CS) parameters: body weight loss, body temperature loss, general behavior, and morbidity signs (appearance and breathing). At the end of the experiments, mice were euthanized with 150mg/Kg sodium pentobarbital (CHUV Pharmacie, Lausanne, Switzerland) to collect samples

### Lung histology

Resected murine lung tissue was fixed in 5mL of 10% buffered formalin and embedded in paraffin. Tissue sections of the left lung were prepared at (i) 4μm and (ii) 6μm and were consequently stained with (i) Grocott Methenamine Silver (GMS) staining and (ii) Hematoxylin and Eosin (H&E) by the Mouse Pathology Facility (UNIL, Lausanne, Switzerland). The slides were analyzed using a Leica DM4B microscope equipped with a Leica DMC2900 camera with Leica Application Suite X (LSX) software version 2.0.0.14332.2.

### Statistical analyses

Statistical significance of the data from the confrontations on differentiated bronchial tissues in Transwell® inserts (3 replicates, n = 3) was tested with unpaired two-tailed Student t-tests in Microsoft® Excel (Version 16.37). The statistical significance threshold was set to 5%.

## Acknowledgments

We would like to thank Eric Bernasconi, Ilona Palmieri, Nourine Noormamode, Aislinn Estoppey, Saskia Bindschedler, Karen Davenport, for their support in the realization of preliminary experiments. We would like to thank Diego Gonzales and Ted Turlings for critical review of the paper.

## Author contributions

Conceptualization: AK, MG, SN, CVG, NDU, PJ

Methodology: NDU, AK, JFH, TJ, SN, MP, JDS, NH

Investigation: FP, AT, AS, AB, JP, TJ, JMK, AV

Visualization: FP, AT, NDU, TJ, SN, PJ

Funding acquisition: MP; AK, PSC, PJ

Project administration: NDU, PJ

Supervision: PJ, NDU, SM

Writing – original draft: FP, TJ, NDU, PJ

Writing – review & editing: All authors

## Data and materials availability

All data are available in the main text or the supplementary materials.

