## Supplementary materials for "Fighting *Aspergillus* infection using biocontrol bacteria: A proof-of-concept of environmental interference in a translational setting"

**Affiliations:**
[1]Laboratory of Microbiology, University of Neuchâtel, Neuchâtel, 2000, Switzerland.
[2]Division of Pulmonary Medicine, Lausanne University Hospital (CHUV) and University of Lausanne, Lausanne, 1011, Switzerland
[3]Vital-IT Group, SIB Swiss Institute of Bioinformatics, Bâtiment Amphipôle, Quartier UNIL-Sorge, 1015 Lausanne, Switzerland
[4]Bioscience Division, Los Alamos National Laboratory, Los Alamos, NM, USA
[5]Neuchâtel Platform of Analytical Chemistry, University of Neuchâtel, Neuchâtel, 2000, Switzerland
[6]Alveolix, Swiss Organs-on-Chip Innovation, 3010, Bern, Switzerland.

**List of Supplementary Materials**

**Table S1. Species composition of the clusters depicted in Figure 1.** The Aspergillus spp. Composing each cluster is depicted in the column "Species". For each species the proportion of clinical (prop_clin) and oxalotrophic (prop_oxal) is provided as well.

**Fig. S1. Effect of increasing *A. niger* (An M8) conidial load (10-10000) on the morphology of bronchial epithelial cells.** From top row to the last: control cells; 10, 50, 100, 500, 5000, and 10000 conidia of An M8. After 24 h incubation, an increase in cell damage with increasing conidial load was observed. Damage was visible from a conidial load of 500. Damaged cells began to shrink, and actin got more agglomerated, compared to the cells-only control. From a conidial load of 1000 and on, fungal growth has an adverse effect on tissue integrity. Visible hyphae are shown by white arrows Culture medium volume was 200 µl/well. Scale bars = 200 µm

**Fig. S2. Effect of increasing *C. oxalaticus* (Co) cell load (10-1000) on the morphology of bronchial epithelial cells.** From top row to the last: control cells; 10, 50, 100, 500, and 1000 bacterial cells of Co. After 24h incubation, an increase in cell damage with increasing bacterial cell load was observed. As few as ten bacterial cells already had an impact on cell morphology. Indeed, cells became rounder, and actin got more agglomerated, compared to the cells-only control. However, the cytopathic effect observed in the presence of An M8 was less pronounced (Figure SXX). Culture medium volume was 200 µl/well. Scale bars = 200 µm.

**Fig. S3. Effect of co-culturing *C. oxalaticus* (Co) with *A. niger* (An M8) on cell morphology in submerged undifferentiated bronchial epithelium.** From top row to the last: control cells; 10 conidia of An M8; 500 conidia; 10 bacterial cells of Co; 10 conidia and 10 bacterial cells; 500 conidia and 10 bacterial cells. After 72h incubation, the damage and cytopathic effect of An M8 conidia was clearly visible with as few as ten conidia per well. The cells appear even more damaged with a conidial load of 500. Ten Co cells also changed the morphology of the epithelial cells, but no cytopathic effect was observed. With the co-inoculation of as few as ten Co cells, the morphology of bronchial cells infected with An M8 was like the morphology of bacteria-only control. Culture medium volume was 200 µl/well. Scale bars = 200 µm.

**Fig. S4. *In vitro* model of the "environmental interference concept" confronting the oxalotrophic bacteria *Cupriavidus oxalaticus* with *Aspergillus niger.*** Representative immunofluorescence microscopy pictures of AECs. (magnification 10x, scale = 100 µm).

**Fig. S5. Pro-inflammatory effect of *C. oxalaticus*.** Using the A549 Luciferase-reporter cells for the induction of inflammation (NF-kB pathway), *C. oxalaticus* was shown to be pro-inflammatory, as shown by the higher level of induction compared to cells alone, as well as when it was co-added with TNFa (1 and 10 ng/mL).

**Fig. S6. Cytotoxicity and epithelial barrier damages by *C. oxalaticus*.** To assess cytotoxicity and epithelial damages caused by *C. oxalaticus*, 10 cells of *C. oxalaticus* were inoculated on differentiated primary bronchial epithelial cell cultures on Transwell inserts. **(A)** Cytotoxicity of *C. oxalaticus* (Co) was assessed through the quantification of LDH levels in the culture supernatant. Inoculation of *C. oxalaticus* cells on bronchial epithelial cell cultures (C + Co) induced significant LDH leakage as compared to cells alone (C). **(B)** Moreover, *C. oxalaticus* induced epithelial barrier damage as shown by higher permeability to Lucifer Yellow (LY) compared to the baseline permeability of the control cells. **(C)** Immunofluorescence picture showing healthy control cells with actin in green, the tight-junction Zona-Occludens 1 ZO-1 protein in red, and the nuclei in blue. **(D)** Immunofluorescence picture showing cells stimulated with 10 *C. oxalaticus*. Actin appears in green, nuclei in blue, and *C. oxalaticus* cells in red. **(E)**. Immunofluorescence pictures were taken with a confocal microscope.

**Figure S7. Kaplan-Meier survival plot of *G. mellonella* larvae injected with C. albicans and S. aureus.** $10^5$ *C. albicans* (Ca) and 2 x $10^4$ *S. aureus* (Sa) were injected both alone and as a co-injection in *Galleria* larvae. Survival was monitored for 72 h.

**Table S1. Species composition of the clusters depicted in Figure 1.** The Aspergillus spp. Composing each cluster is depicted in the column "Species". For each species the proportion of clinical (prop_clin) and oxalotrophic (prop_oxal) is provided as well.

| **Cluster** | **Species** | **prop_clin** | **prop_oxal** |
|---|---|---|---|
| brasiliensis | *brasiliensis* | 0 | 0.67 |
| flavus | *flavus* | 0.1 | 1 |
| fumigatus | *fumigatus* | 0.24 | 1 |
| jensenii | *hiratsukae* | 1 | 0 |
| jensenii | *jensenii* | 1 | 0 |
| jensenii | *nidulans* | 1 | 0 |
| jensenii | *spinulosporus* | 1 | 0 |
| jensenii | *turcosus* | 1 | 0 |
| latus | *latus* | 1 | 0.17 |
| niger | *niger* | 0.2 | 0.98 |
| oryzae | *aff.* | 0 | 1 |
| oryzae | *affinis* | 0 | 1 |
| oryzae | *bridgeri* | 0 | 1 |
| oryzae | *brunneus* | 0 | 1 |
| oryzae | *burnettii* | 0 | 1 |
| oryzae | *cejpii* | 0 | 1 |
| oryzae | *cristatus* | 0 | 1 |
| oryzae | *diversus* | 0 | 1 |
| oryzae | *elegans* | 0 | 1 |
| oryzae | *fischeri* | 0 | 1 |
| oryzae | *funiculosus* | 0 | 1 |
| oryzae | *haitiensis* | 0 | 1 |
| oryzae | *hancockii* | 0 | 1 |
| oryzae | *intermedius* | 0 | 1 |
| oryzae | *minisclerotigenes* | 0 | 1 |
| oryzae | *montevidensis* | 0 | 1 |
| oryzae | *nomiae* | 0 | 1 |
| oryzae | *ochraceus* | 0 | 1 |
| oryzae | *olivimuriae* | 0 | 1 |
| oryzae | *oryzae* | 0 | 1 |
| oryzae | *pseudoglaucus* | 0 | 1 |
| oryzae | *sclerotialis* | 0 | 1 |
| oryzae | *sclerotiorum* | 0 | 1 |
| oryzae | *sojae* | 0 | 1 |

| | | | |
|---|---|---|---|
| oryzae | *sparsus* | 0 | 1 |
| oryzae | *texensis* | 0 | 1 |
| oryzae | *viridinutans* | 0 | 1 |
| oryzae | *westerdijkiae* | 0 | 1 |
| parasiticus | *parasiticus* | 0 | 0.8 |
| tamarii | *tamarii* | 0.5 | 1 |
| terreus | *terreus* | 0.17 | 1 |
| thermomutatus | *felis* | 1 | 1 |
| thermomutatus | *hortae* | 1 | 1 |
| thermomutatus | *thermomutatus* | 1 | 1 |
| tubingensis | *tubingensis* | 0.25 | 0.17 |
| ustus | *aculeatinus* | 0 | 0 |
| ustus | *amoenus* | 0 | 0 |
| ustus | *asperescens* | 0 | 0 |
| ustus | *baeticus* | 0 | 0 |
| ustus | *calidoustus* | 0 | 0 |
| ustus | *cavernicola* | 0 | 0 |
| ustus | *conjunctus* | 0 | 0 |
| ustus | *costaricensis* | 0 | 0 |
| ustus | *croceus* | 0 | 0 |
| ustus | *fijiensis* | 0 | 0 |
| ustus | *incahuasiensis* | 0 | 0 |
| ustus | *japonicus* | 0 | 0 |
| ustus | *kassunensis* | 0 | 0 |
| ustus | *luchuensis* | 0 | 0 |
| ustus | *nanangensis* | 0 | 0 |
| ustus | *ochraceoroseus* | 0 | 0 |
| ustus | *panamensis* | 0 | 0 |
| ustus | *puulaauensis* | 0 | 0 |
| ustus | *quadricinctus* | 0 | 0 |
| ustus | *quadrilineatus* | 0 | 0 |
| ustus | *rambellii* | 0 | 0 |
| ustus | *sydowii* | 0 | 0 |
| ustus | *unguis* | 0 | 0 |
| ustus | *ustus* | 0 | 0 |
| ustus | *versicolor* | 0 | 0 |
| welwitschiae | *welwitschiae* | 0.25 | 1 |

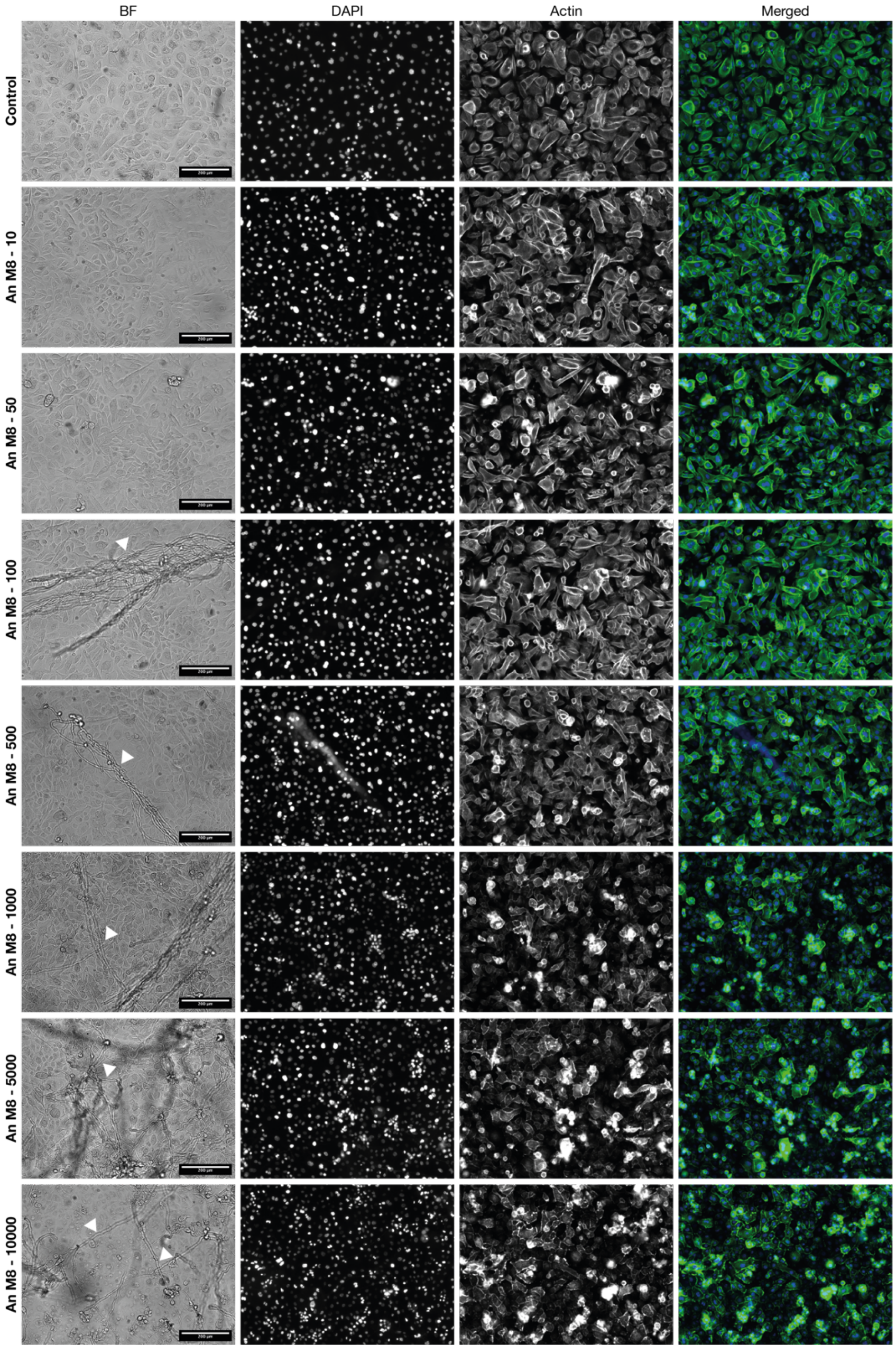


**Fig. S1. Effect of increasing *A. niger* (An M8) conidial load (10-10000) on the morphology of bronchial epithelial cells.** From top row to the last: control cells; 10, 50, 100, 500, 5000, and 10000 conidia of An M8. After 24 h incubation, an increase in cell damage with increasing conidial load was observed. Damage was visible from a conidial load of 500. Damaged cells began to shrink, and actin got more agglomerated, compared to the cells-only control. From a conidial load of 1000 and on, fungal growth has an adverse effect on tissue integrity. Visible hyphae are shown by white arrows Culture medium volume was 200 µl/well. Scale bars = 200 µm.

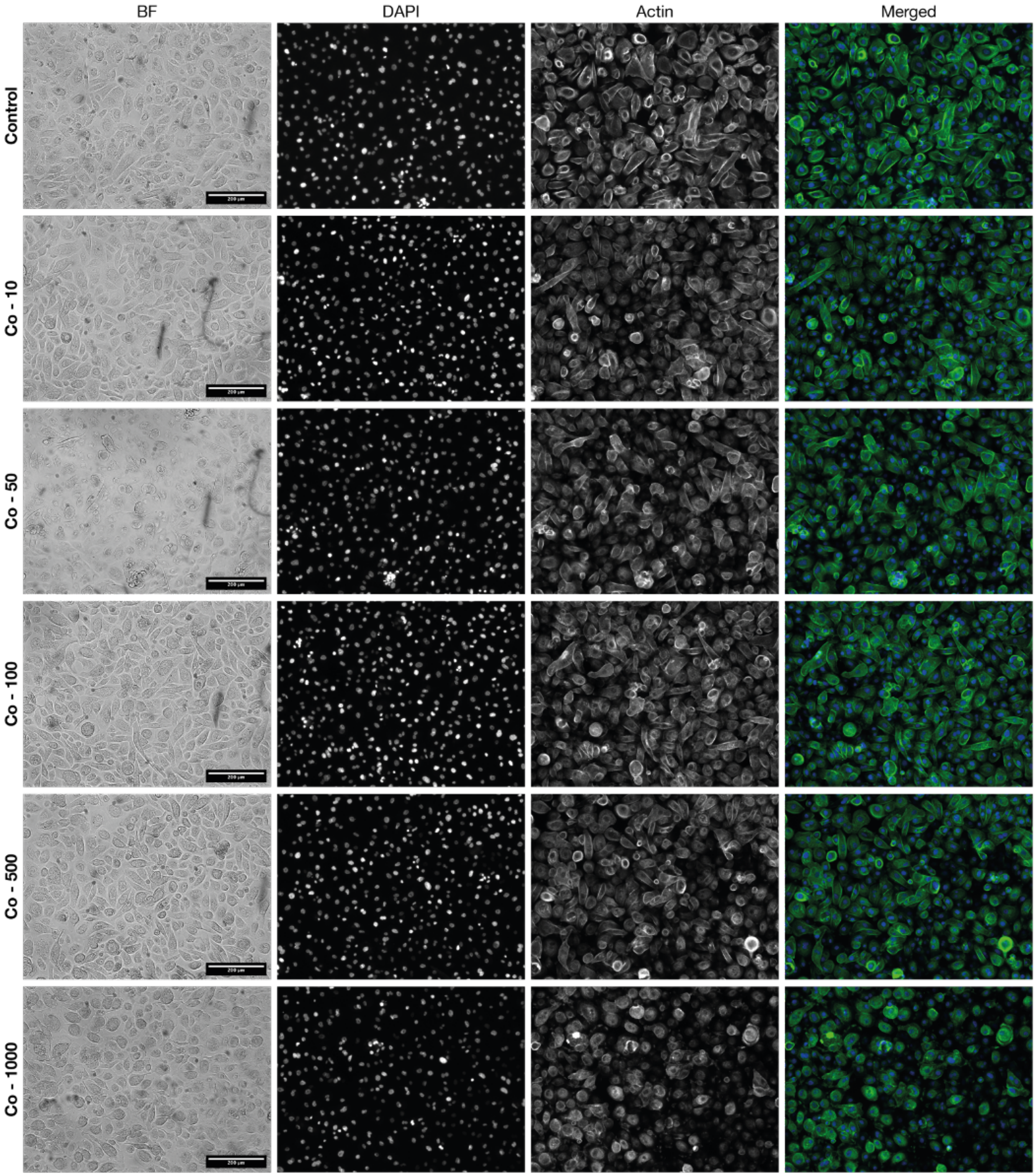


**Fig. S2. Effect of increasing *C. oxalaticus* (Co) cell load (10-1000) on the morphology of bronchial epithelial cells.** From top row to the last: control cells; 10, 50, 100, 500, and 1000 bacterial cells of Co. After 24h incubation, an increase in cell damage with increasing bacterial cell load was observed. As few as ten bacterial cells already had an impact on cell morphology. Indeed, cells became rounder, and actin got more agglomerated, compared to the cells-only control. However, the cytopathic effect observed in the presence of An M8 was less pronounced (Figure SXX). Culture medium volume was 200 µl/well. Scale bars = 200 µm.

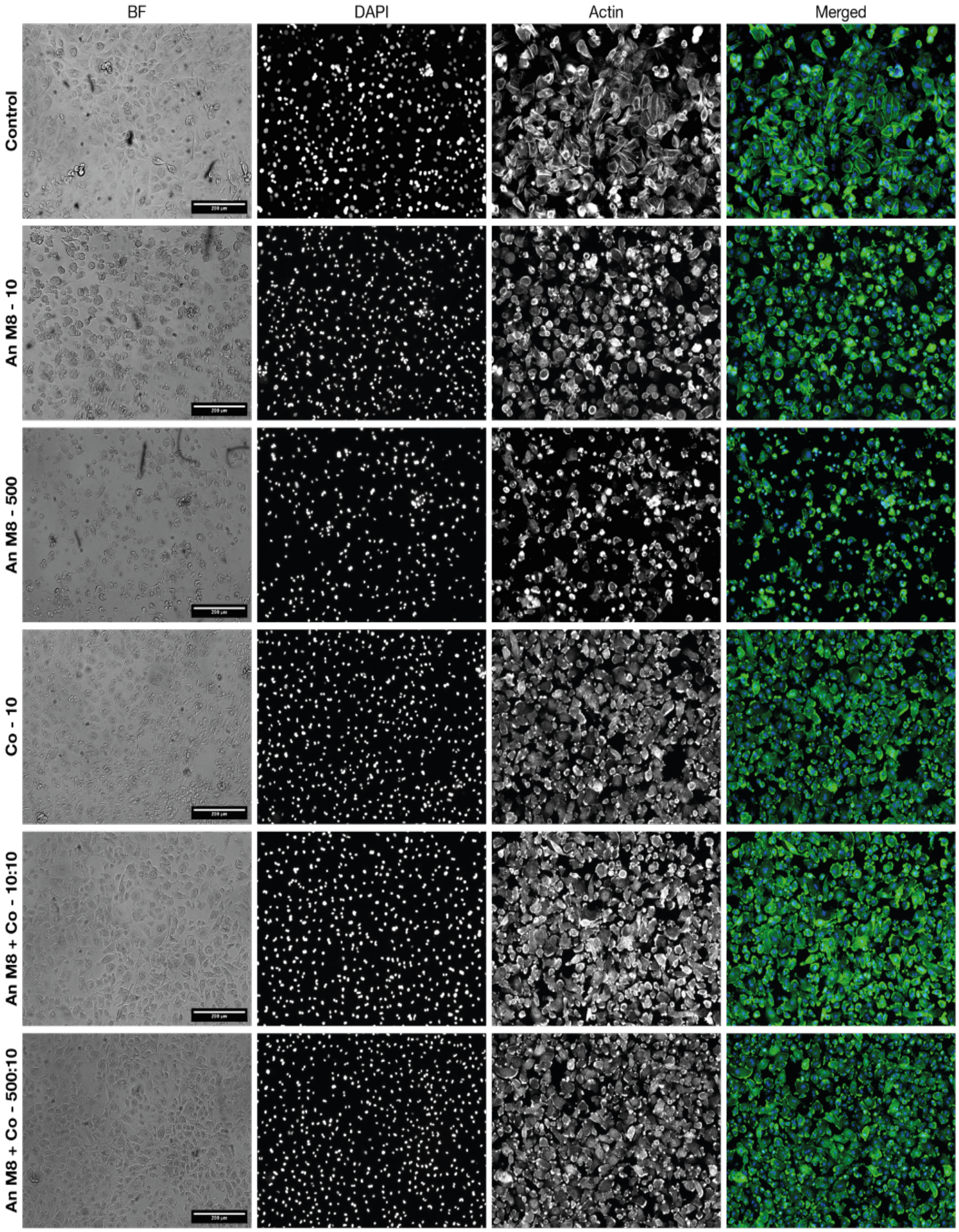


**Fig. S3. Effect of co-culturing *C. oxalaticus* (Co) with *A. niger* (An M8) on cell morphology in submerged undifferentiated bronchial epithelium.** From top row to the last: control cells; 10 conidia of An M8; 500 conidia; 10 bacterial cells of Co; 10 conidia and 10 bacterial cells; 500 conidia and 10 bacterial cells. After 72h incubation, the damage and cytopathic effect of An M8 conidia was clearly visible with as few as ten conidia per well. The cells appear even more damaged with a conidial load of 500. Ten Co cells also changed the morphology of the epithelial cells, but no cytopathic effect was observed. With the co-inoculation of as few as ten Co cells, the morphology of bronchial cells infected with An M8 was like the morphology of bacteria-only control. Culture medium volume was 200 µl/well. Scale bars = 200 µm.

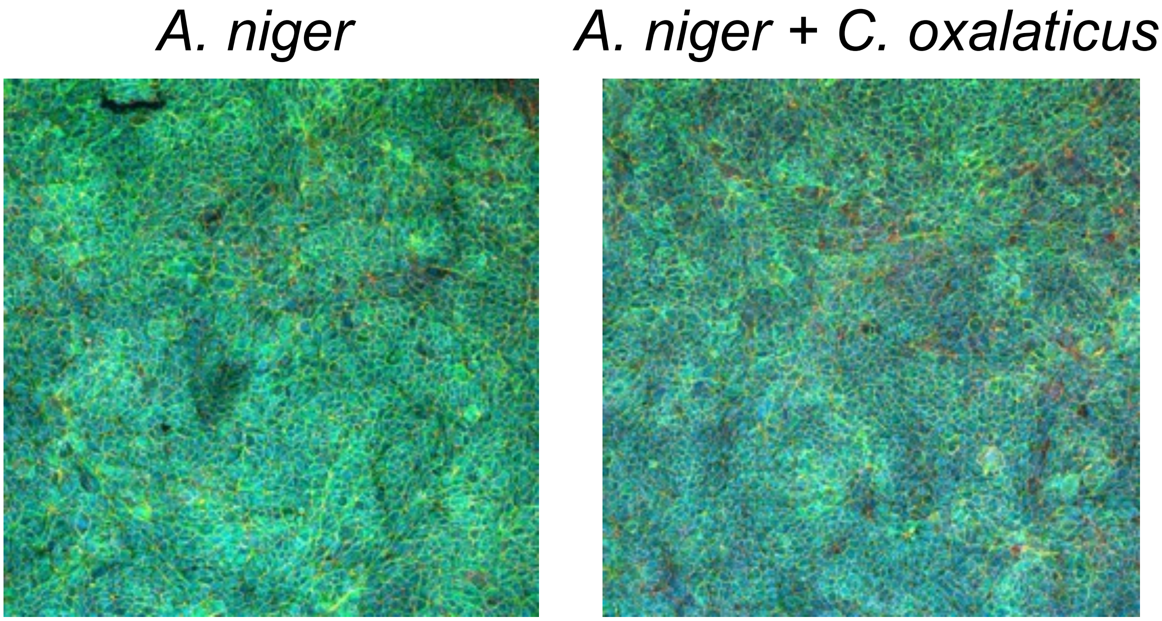


**Fig. S4. *In vitro* model of the "environmental interference concept" confronting the oxalotrophic bacteria *Cupriavidus oxalaticus* with *Aspergillus niger*.** Representative immunofluorescence microscopy pictures of AECs. (magnification 10x, scale = 100 µm).

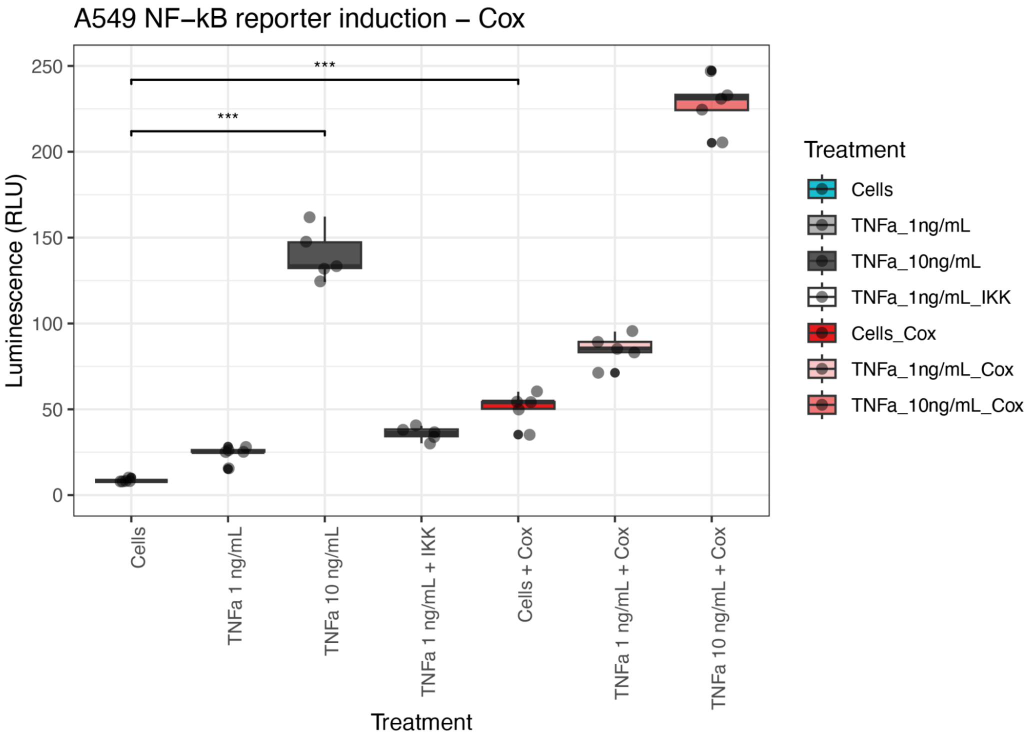


**Fig. S5. Pro-inflammatory effect of *C. oxalaticus*.** Using the A549 Luciferase-reporter cells for the induction of inflammation (NF-kB pathway), *C. oxalaticus* was shown to be pro-inflammatory, as shown by the higher level of induction compared to cells alone, as well as when it was co-added with TNFa (1 and 10 ng/mL).

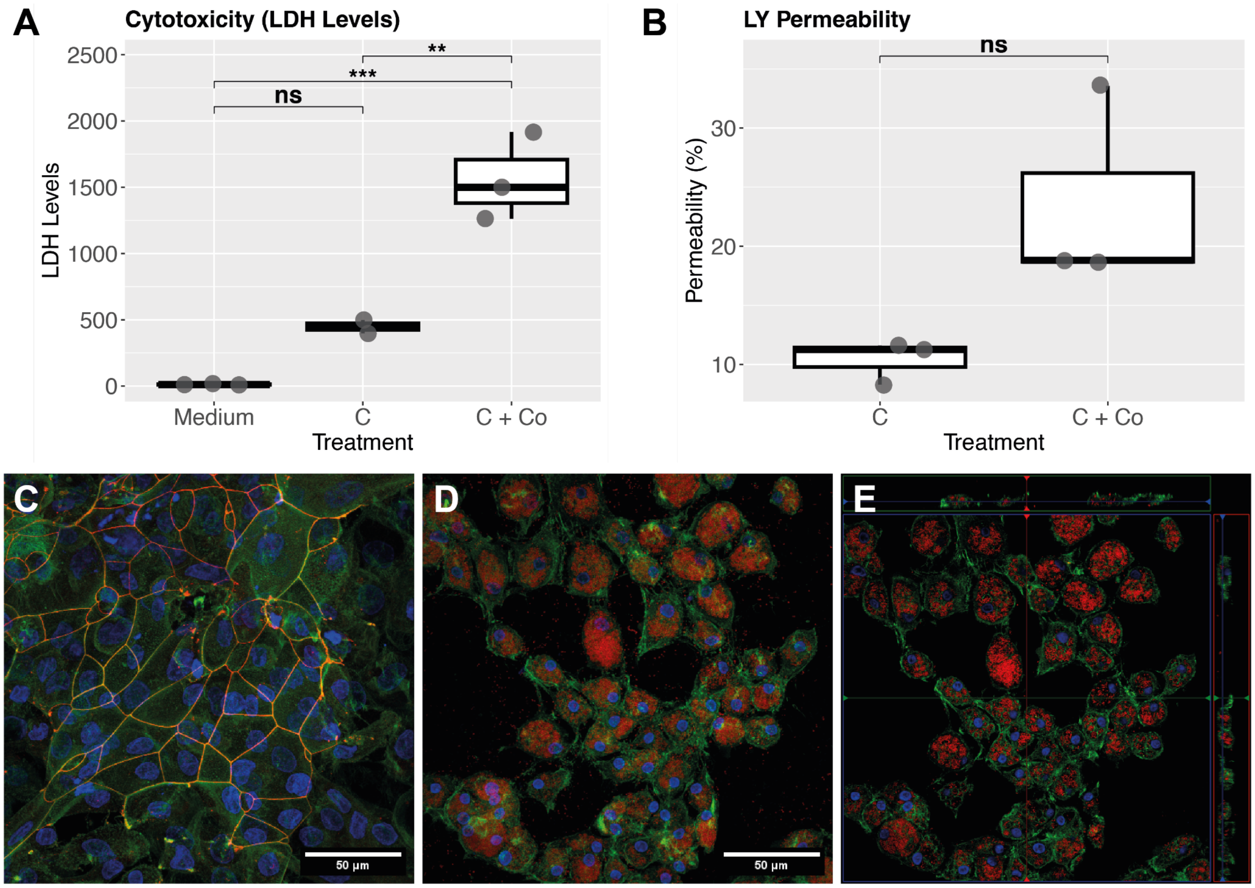


**Fig. S6. Cytotoxicity and epithelial barrier damages by *C. oxalaticus*.** To assess cytotoxicity and epithelial damages caused by *C. oxalaticus*, 10 cells of *C. oxalaticus* were inoculated on differentiated primary bronchial epithelial cell cultures on Transwell inserts. **(A)** Cytotoxicity of *C. oxalaticus* (Co) was assessed through the quantification of LDH levels in the culture supernatant. Inoculation of *C. oxalaticus* cells on bronchial epithelial cell cultures (C + Co) induced significant LDH leakage as compared to cells alone (C). **(B)** Moreover, *C. oxalaticus* induced epithelial barrier damage as shown by higher permeability to Lucifer Yellow (LY) compared to the baseline permeability of the control cells. **(C)** Immunofluorescence picture showing healthy control cells with actin in green, the tight-junction Zona-Occludens 1 ZO-1 protein in red, and the nuclei in blue. **(D)** Immunofluorescence picture showing cells stimulated with 10 *C. oxalaticus*. Actin appears in green, nuclei in blue, and *C. oxalaticus* cells in red. **(E)**. Immunofluorescence pictures were taken with a confocal microscope.

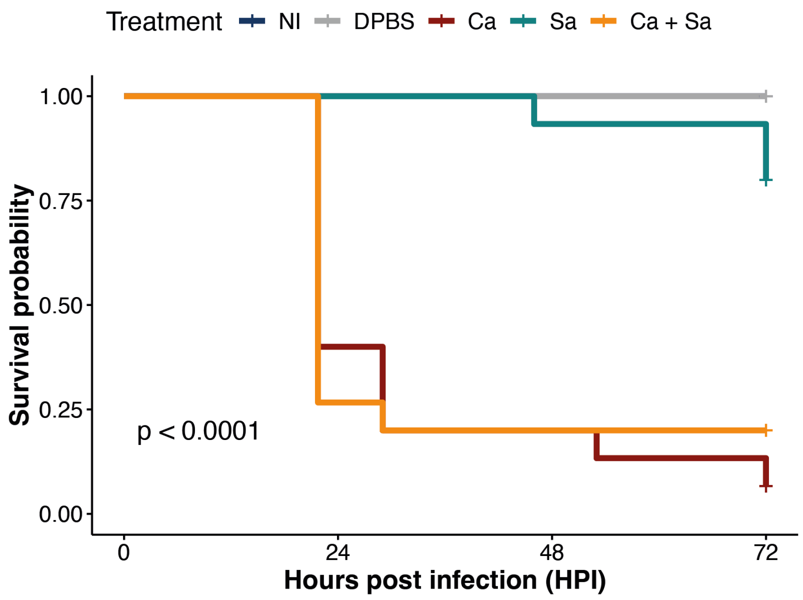


**Figure S7. Kaplan-Meier survival plot of *G. mellonella* larvae injected with C. albicans and S. aureus.** $10^5$ *C. albicans* (Ca) and 2 x $10^4$ *S. aureus* (Sa) were injected both alone and as a co-injection in *Galleria* larvae. Survival was monitored for 72 h.
